# A Python Toolbox for Representational Similarity Analysis

**DOI:** 10.1101/2025.05.22.655542

**Authors:** Jasper J.F. van den Bosch, Tal Golan, Benjamin Peters, JohnMark Taylor, Mahdiyar Shahbazi, Baihan Lin, Ian Charest, Jörn Diedrichsen, Nikolaus Kriegeskorte, Marieke Mur, Heiko H. Schütt

## Abstract

Representational similarity analysis (RSA) is a method to characterize neural representations and evaluate computational models based on neural representational geometries. Here we present a wave of recent methodological advances, including improved measures of representational distances, evaluators for representational models, and statistical inference methods, which are available to the community in a new open-source toolbox in Python. The rsatoolbox enables neuroscientists to explore neural representational geometries and to evaluate neural network models, connecting theory to experiment in the new era of big models and big data.

## 1 Introduction

Neuroscience now has many methods to measure brain activity with ever increasing precision and coverage. Recently, these improved methods have begun to be used to collect large datasets of brain responses to natural stimuli [e.g. 1–6]. These large datasets aim to constrain the complex computational models we need to capture the complexity of brain processing in our complex world [7–9]. Recent advances in deep learning and AI [10–13] now allows to develop complex models that can perform cognitive tasks at scale. To connect our complex models to the large datasets poses a new set of statistical challenges. Here we present an integrated methodology implemented in an open-source Python toolbox that addresses these challenges.

The evaluation of models with neural activity measurements is usually based on comparisons of internal representations, i.e. whether internal activities computed from the input are similar. The data for such comparisons are matrices of activities with dimensions for the conditions and the measurement channels (fMRI Voxels, MEG/EEG channels, neurons, electrodes/multiunit activities, features in a model, etc.). We can compare these activity matrices, whether or not our brains and/or models represent the condition in a meaningful way. Representing something would imply that these activity patterns are used for some computation [14–16], or at very least provide information about the conditions.

We focus on a particular method for comparing representations called representational similarity analysis [RSA 17]. See section 3.3 for alternative approaches. RSA compares models or brains based on the dissimilarities between the response patterns elicited by the different conditions. These dissimilarities determine how well a downstream area could decode which condition was presented and the geometry of the activity patterns [18]. Critically, RSA allows us to compare activity matrices without creating a mapping between features and response channels. In practice, RSA entails four steps: First, we pre-process data to estimate the activity pattern for each stimulus or condition. Second, we compute the dissimilarity between each pair of conditions for both the data and the model. A matrix of dissimilarity values across all conditions is called a Representational Dissimilarity Matrix (RDM). Third, we compare the RDMs. Fourth, we perform statistical inference to estimate the uncertainty about our results. RSA is historically rooted in the use of multidimensional scaling (MDS) to infer the geometry of mental representations on the basis of behavioral data [19, 20] and in the concept of second-order isomorphism, which posits that relationships among cognitive representations are isomorphic to relationships among the things in the world that they represent [21, 22].

RSA has enabled numerous and impactful research studies. The abstraction from measurement channels inherent to RSA has allowed us to compare brains across species [18, 23] and age groups [24]; across measurement modalities [25–27], and across stimulus modalities (e.g. orthographic and visual [28], orthographic and phonological [29], natural sounds [30]). RSA has been employed across many fields of study as diverse as linguistics [31] anthropology [32] and computational psychiatry [33], but particularly in neuroscience to study perception [e.g. 34–37] and memory [38–40]. Although RSA has predominantly been popular in human neuroimaging, there is a growing body of literature with RSA on neural recordings [41–46]. Another promising avenue is that of relating Artificial Neural Network models to brain data using RSA [25, 44, 47–57], but see [58]. RSA has been related to other analysis perspectives such as neural tuning curves [59], linear encoding models [60, 61], generalized shape metrics [62], kernel based methods [63], and pattern component modeling [64].

Since its inception many improvements were added to RSA, which make the method more powerful, but require more complex implementations. First, statistical considerations [65, 66] provide strong arguments in favor of other measures of dissimilarity for the construction of RDMs than the classical correlation distance [17]. In particular, (squared) euclidean distances, Mahalanobis distances that take noise covariances into account, and cross-validated distances that remove the positive bias contained in distances computed from noisy signals are now standard [65]. Second, new metrics have been proposed for comparing RDMs: It has been proposed to use a distance on the Riemannian manifold of positive semi-definite matrices [67]. It has been proposed to use transformations of the distances that focus the analysis more on topology [68, 69]. Finally, whitened cosine similarity and correlation were proposed to compensate for the correlations between distances in the RDM [70]. Third, the methodology for statistical inference in RSA has crystallized: In RSA, we usually want to generalize over conditions as well as subjects, which requires complex bootstrap methods for accurate inference [71]. Additionally, model evaluation for flexible models is now possible by using cross-validation techniques [72]. Although the basic principles of RSA are straightforward to implement, many of the improvements to RSA are not. Thus, a wider adoption of the improvements requires a standardized and validated implementation of the best practices for RSA.

Here, we present our new *rsatoolbox*, which fills this gap with an implementation in Python that includes the new developments. We choose Python [73], because the neuroscience community has adopted Python as a programming language of choice. Python’s open-source nature and the batteries-included philosophy have led to a number of key tools, like NiBabel [74], the NIPY ecosystem [75], MNE [76], fMRIprep [77], or Neurodata without borders [NWB; 78–80] and others [81–83]. Therefore, it makes sense for our new library to plug into this growing ecosystem of Python tools for neuroscientists.

The body of this manuscript follows the workflow of a typical RSA analysis with new developments, their validation, best practice recommendations and a guide for how these can be performed using the rsatoolbox with examples. Afterwards we discuss the current state and outlook on how the toolbox may be extended in the future, integration into the wider context of Python packages for neuroscience and compare to alternative approaches for comparisons of brains and models.

## 2 Representational similarity analysis step by step

### Step 1: Importing data and estimating activity patterns

Data to be analyzed with RSA varies widely in its dimensions and how it is annotated. To bridge the gap between these varied data formats, the first analysis step is to bring the raw data into a common input structure, the *Dataset* class (see Fig. 2 for an overview). This class then provides methods for selection and organization and serves as a standardized input format for the rest of the toolbox.

**Fig. 1.**
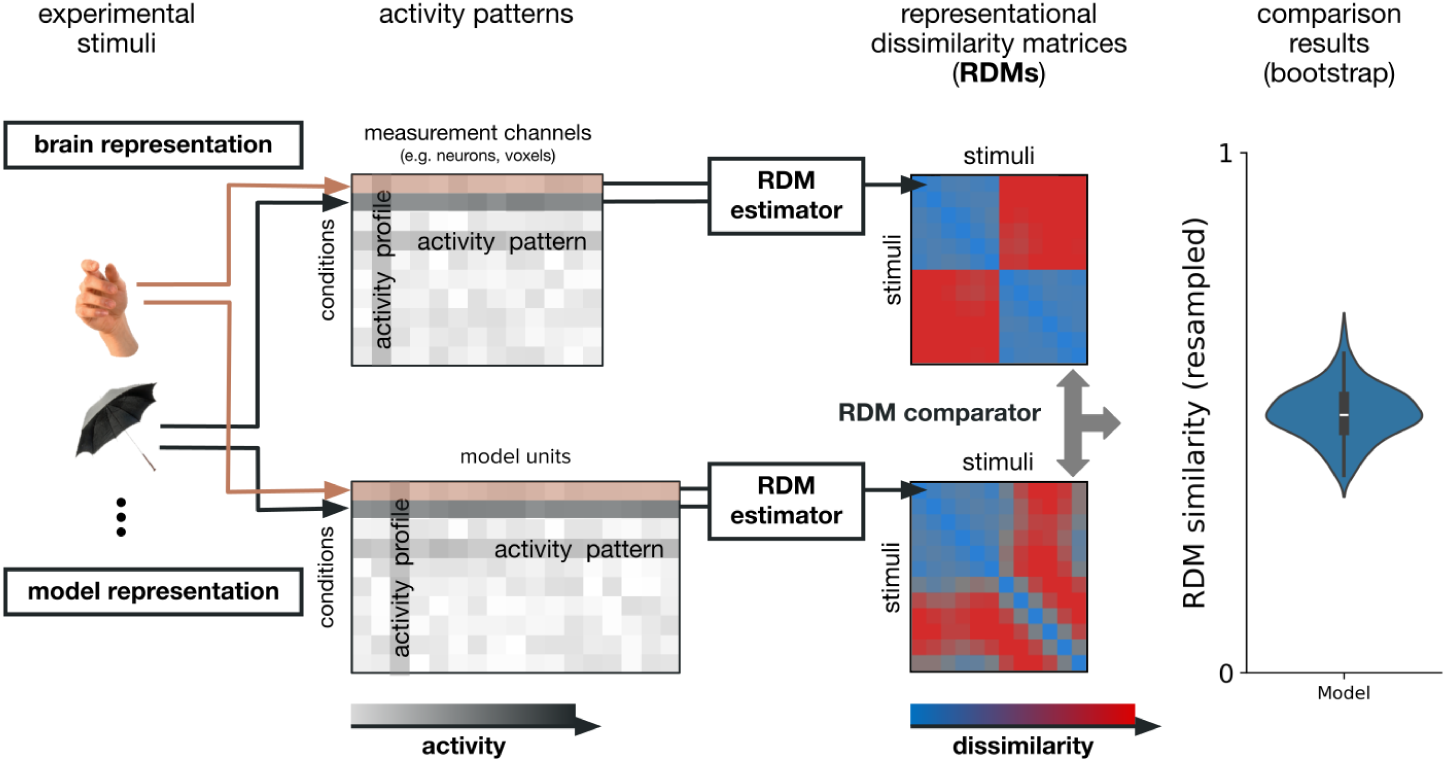
Processing flow for representational similarity analysis (RSA). Experimental stimuli or conditions are presented to both the subjects and the model we want to test. Using an estimator for the representational dissimilarity matrix (RDM), we compute the dissimilarities between all pairs of stimuli. These matrices can then be compared using an RDM comparator. Uncertainty estimates for the comparison results can be obtained from bootstrap resampling.

**Fig. 2.**
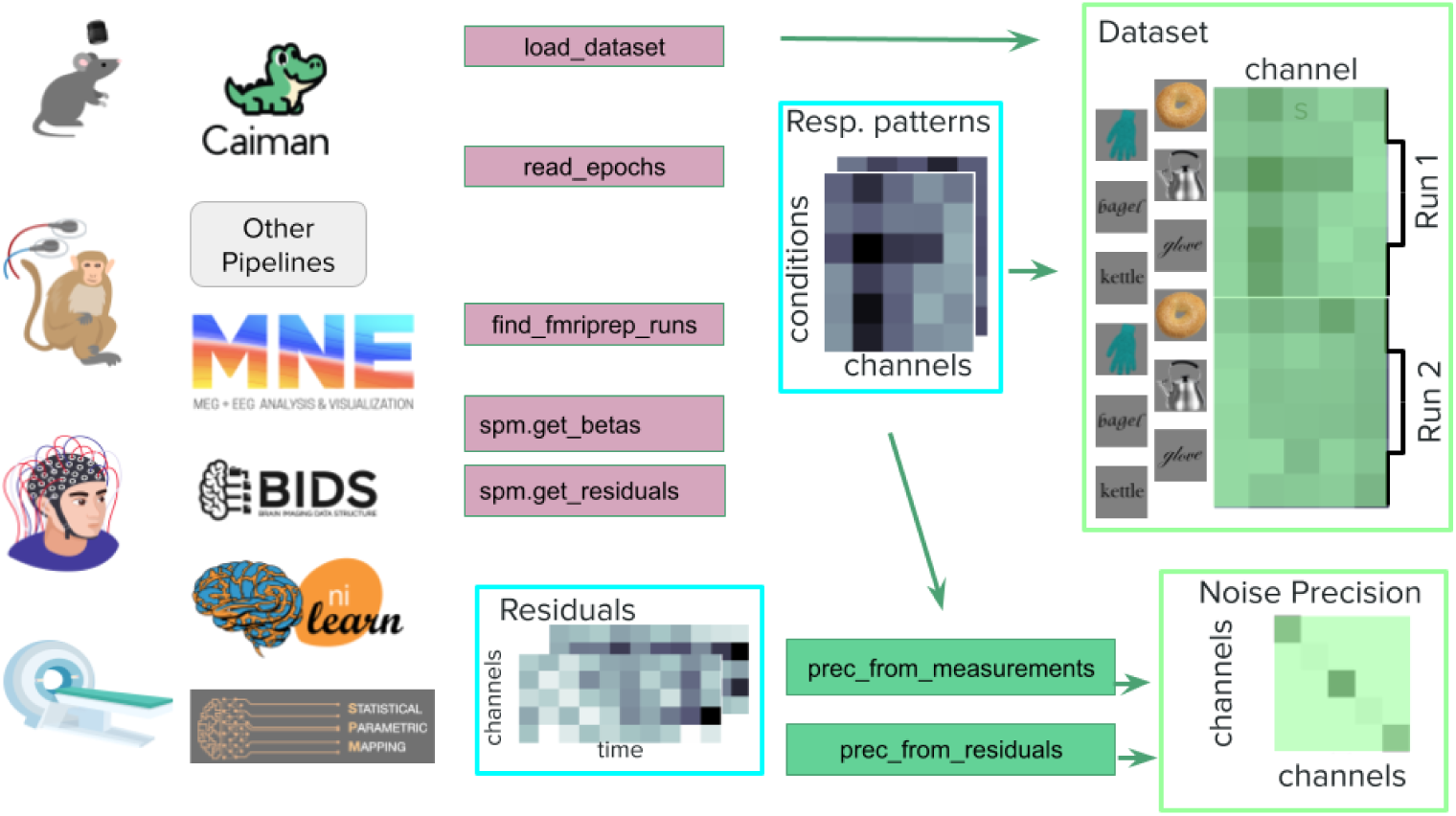
Preprocessing workflow. RSA can be applied to data from a wide variety of sources. Neural data recorded from any species, in any modality such as single-cell recordings, calcium imaging, magnetoencephalography (MEG), electroencephalography (EEG), local field potentials (LFP), and functional magnetic resonance imaging (fMRI). RSAtoolbox integrates with a range of popular analysis tools to help prepare and import data patterns as well as residuals. This is in the form of i/o functions to read various data formats as well as utilities to make it easier to navigate a dataset. For examples applied to fMRI data, see the SPM demo, the nilearn demo, or a bare-bones NumPy patterns demo.

The main content of a Dataset object is a matrix of activity data, with rows corresponding to individual measurements and columns to measurement channels like voxels, sensors, or neurons. Additionally, the object contains annotations, including the condition each measurement corresponds to and optional additional annotations about the measurements and channels. To use cross-validated distance estimates (see section 2.3), multiple independent estimates for the activity pattern for each condition are necessary. To enable this approach, the Dataset class can contain multiple measurements for the same condition. While Dataset objects can be created simply from a NumPy array and lists of annotations, we provide some modality-specific functions that assist in importing data and processing it in a manner optimized for RSA for ease of use.

Additional to the response strengths, some analyses require an estimate of the signal covariance across measurement channels, i.e. of ‘noise correlations‘. There are two main reasons to take this covariance into account: First, it aligns the distance estimates better with statistical discriminability (as discussed more below). Second, it can improve the reliability of distance estimates because channels with high variance or strong correlations with other channels are given lower weights for the computation of distances [65].

Given that different measurement modalities require different steps for estimating activity patterns and their noise covariance, we briefly provide separate recommendations for neural spiking data, fast averaged data like EEG, MEG or local field potentials, fMRI data, and behavioral data.

#### Neural recordings/spiking data

For neural data, we aim to extract a spike rate for each neuron and each condition. The simplest method to do so is to count spikes in a fixed time window after stimulus presentation. It is possible to add further preprocessing like temporal smoothing, artifact removal or similar adjustments as they are deemed necessary for the specific brain area and recording technique. We generally advise against subtracting baselines, because they regularly result in negative spike rates which breaks a central assumption of many methods adapted to spike rate distributions. The main exception are brain areas with strong, slow drifts of the average spike rate, whose effect can be substantially reduced by subtracting an individual baseline for each trial.

Spike rates are typically more variable if the mean response is higher, i.e. they are not homoscedastic. The classic assumption is Poisson noise, which has variance proportional to the mean response [84]. More modern takes acknowledge that the rates can be more or less variable than expected from a Poisson distribution and follow more complex distributions in general [85–88]. The standard Euclidean and Mahalanobis distances are thus a bad fit for spike rate data. We provide two approaches to this problem: First, the spike rates can be transformed such that they are better approximated by a homoscedastic noise model. The most common transformation is a square root transform [89]. Second, we can use different distance measures that take the difference in variance into account. For this purpose we provide the symmetrized KL-divergence between Poisson distributions (see below). For this approach, the spike rates are not transformed.

Noise correlation between neurons is weaker and less problematic than noise correlation between channels in other modalities; therefore, RSA on spiking data does not strictly require taking the noise covariance matrix into account. If this is desired nonetheless, the estimate for the covariance is usually based on the variance of individual responses around the mean response for each condition. This can be computed from the Dataset object in the toolbox, such that no separate noise estimate needs to be extracted unless you wish to use an alternative noise covariance estimate.

When the temporal evolution of processing is of interest, all processing in the RSA toolbox can be repeated for different time points. To do so, we simply compute datasets for each time point we want to test by adjusting the window we count spikes over or taking a different time point from a kernel density estimate of spike rate. This can be done at equal time steps after stimulus onset to get a curve of results over time. To enable this the rsatoolbox provides a *TemporalDataset* class. Alternatively, the analysis can be focused on particular phases of the response, for example the initial onset response and the sustained activity. The latter approach typically results in much fewer, more targeted analyses.

#### Electroencephalography and Magnetoencephalography

For Electroencephalography (EEG), Magnetoencephalography (MEG), or their combination (M/EEG), RSA analyses are based on the average responses at the sensor or source level to the respective conditions. Using standard toolboxes (e.g., MNE-Python, [76]) we first preprocess data (filtering, artifact removal, etc.), optionally project it into source space, and segment it into epochs time-locked to the stimulus onset. We can then estimate pattern response strengths for the different conditions as averages over epochs.

Noise covariance estimates for EEG or MEG can be critical, because both sensor measurements and source estimates exhibit strong spatial correlations. Sensors may also vary substantially in their sensitivity and response magnitudes of different sensor types vary by orders of magnitude (i.e., gradiometers, magnetometers, EEG channels). Estimates for the covariances between channels or source level estimates can be obtained from resting state or baseline data or from the deviations of the raw EEG-MEG signals around the mean response to the stimuli. For computing these covariances, standard methods are available, for example in MNE-Python [76, 90, 91]. The noise covariance can also be estimated within the rsatoolbox using the residuals around the condition mean patterns obtained from the epoched data. Which of these estimates performs best is data set dependent. The covariance of the raw signal, especially during a baseline, can be different from the covariance of the average signal during stimulus presentation, but we usually have more data points to estimate the covariance of the raw signal. As for fMRI, it is almost always advisable to regularize the covariance estimates, i.e. to use shrinkage estimates, or to use a diagonal matrix to perform univariate normalization.

The simplest analysis for M/EEG data performs RSA on a single time point or the average within a single time window. For instance, this approach can be applied to focus the analysis on classical components of event-related potentials. M/EEG data also lends itself well to time-resolved analysis of representational geometries (“RDM movie”). For this purpose, we compute a dataset for each time point we want to test and repeat the analysis on each dataset using the *TemporalDataset* class.

#### Local field potentials

(LFPs) can be handled like EEG or MEG data. After preprocessing the data, the average response at a time point relative to stimulus onset specifies a response strength estimate that is a good basis for an RSA analysis.

#### Functional magnetic resonance imaging

fMRI experiments are usually acquired in different imaging runs, which provide a natural way of breaking up the data set into independent partitions for cross-validation [92]. The standard approach for estimating fMRI activity patterns is a mass-univariate general linear model (first-level GLM), which can be used both for event-related and blocked designs [93]. In a GLM, the response to each condition is usually modeled with a single regressor per imaging run. Recently, there are also methods to estimate beta coefficients for each stimulus presentation instead [94, 95]. Such single trial estimates can be more accurate, but are correlated within a run such that we usually recommend to average them within a run before entering them into the RSA analysis. The resultant regression coefficients (beta-weights) constitute the activity patterns elicited by the condition.

fMRI voxels differ substantially in their signal-to-noise ratio and can be strongly correlated over space. Thus, we recommend to *pre-whiten* the noise of the activity estimates before RSA [65]. The easiest way to do this is to divide the beta-weights of each voxel by the estimate of the noise standard deviation from the GLM (univariate prewhitening). A more advanced approach is to also take into account the noise covariance between voxels (multivariate prewhitening). For computational efficiency reasons, we implement prewhitening as part of the distance calculation as a Mahalanobis distance, i.e. we pass the original beta weights and the covariance matrix to the distance calculation instead of prewhitening the estimates.

To estimate the noise covariance among voxels efficiently, we require access to the full residual time-series from the first-level GLM, which are not saved by default in different neuroimaging packages. Our toolbox therefore contains a series of functions (**rsa.io**) to efficiently import and prewhiten the beta-weights from different neuroimaging packages. These functions account for package-specific GLM-related data structures. Currently, our code supports the import of GLMs performed in Statistical Parametric Mapping [SPM, 96] or in nilearn using the standard naming conventions of BIDS [Brain Imaging Data Structure, 97].

#### Behavioral data

Many behavioral paradigms are suitable for RSA if they can be applied to a varied set of stimuli. Direct similarity judgments are experiments where the participant makes an explicit judgment about the perceived similarity of stimuli. This includes paradigms where a discrete choice is made, such as the same-different task [98], match-to-sample [99], or the odd-one-out task [100]. To convert such data into an RDM, there are two broad approaches: The simpler one is to count the number of choices that speak in favor of a pair of objects being dissimilar from each other. This approach works for judgments about a single dissimilarity and also for triplet tasks were we count how often a pair of stimuli was chosen to be more dissimilar than another one. Typically, this will not result in a valid distance matrix initially, but can be projected onto the cone of valid distance matrices. Another drawback of the simple approach is that it requires (many) judgments for each individual dissimilarity in the matrix. Collecting this amount of data becomes infeasible quickly with growing number of conditions. This motivates the more complex approach of training an embedding model that predicts the behavioural choices [100, 101] These models place each condition on a number of “embedding” dimensions and predict the behavioural choices based on the distances in this embedding space. This approach allows us to fill in distances for which we have no data and additionally yields scores on the embedding dimensions that can be made interpretable.

Other tasks ask observers to judge similarities on continuous scales. The simplest such task is the drag-and-rate task, where observers are asked to place stimulus pairs on a continuous dissimilarity scale [102, 103]. For an overview of direct similarity judgment methods, see [104]. For larger datasets the number of combinations may rule out pair-wise trials; the inverse MDS task or “multiple arrangement” provides an alternative where a larger subset of stimuli are arranged with respect to each other in a single space [105]. Data from many arrangements is pooled by iteratively scaling the partially filled RDMs with respect to the dissimilarity of pairs of items they have in common, and then averaging them. This paradigm has been applied to a number of RSA studies, including those with objects [106], scenes [1], faces [34] and videos [107, 108]. Similarity judgments collected using any of these tasks can be imported directly using the library’s tools for reading table data.

Another approach is to obtain indirect measures of perceptual similarity; where one or several independent variables are measured when the participant perceives the stimuli. One example would be the reaction time in an RSVP task such as the Attentional Blink [109]. In another example, participants wrote sentence captions for displayed images; the corresponding high-dimensional embeddings of which were then used as patterns to calculate a semantic RDM [110]. Others have used mouse tracking patterns [111]. Such indirect data can be imported as Dataset for subsequent calculation of the dissimilarity measure.

### Step 2: Estimating representational geometries

The activity patterns for the experimental conditions form the basis for estimating the representational geometries. The geometry is defined by a distance matrix: a square matrix, containing the distance for each pair of activity patterns. However, different distance measures can be used which vary in the ways they handle (1) the variation of the activity patterns along the population-mean dimension, (2) the correlated noise in the multivariate activity space, and (3) the positive bias associated with naive measurements of distances caused by measurement noise. These three complications are addressed in turn in the next three sub-sections. Our new toolbox offers a wide variety of representational dissimilarity estimators. Table 1 gives an overview of some important representational dissimilarity estimators. An RDM is computed by calling calc_rdm on the Dataset object, with the chosen dissimilarity estimator specified in the string argument method (e.g. method=‘euclidean’, ‘correlation’, ‘mahalanobis’, ‘poisson’, ‘crossnobis’).

**Table 1.**
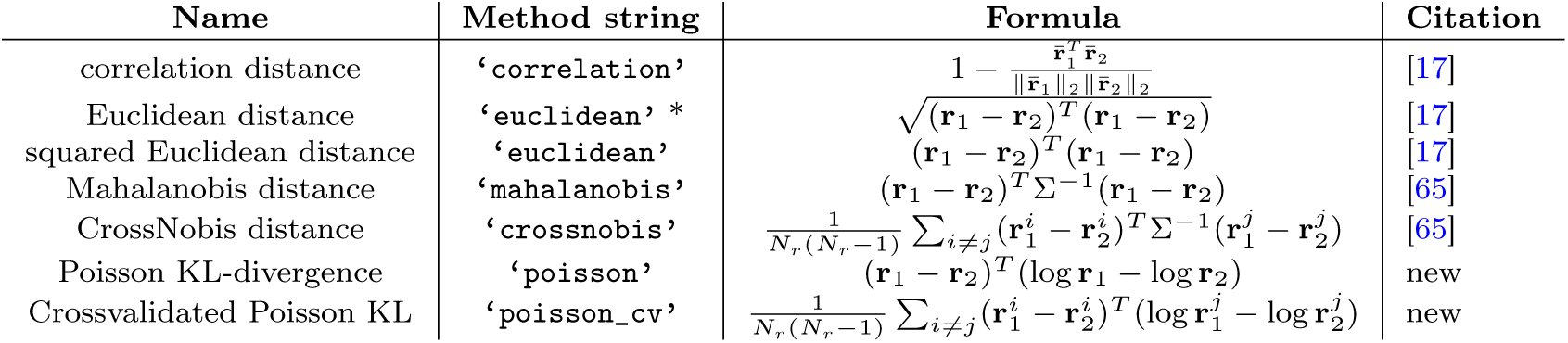
Methods for estimating RDMs. See text for details and recommendations for choosing from this list. **r**_1_ and **r**_2_ are the two response patterns, **r̄**_1_*_/_*_2_ are standardized versions of them, Σ is an estimate of the noise covariance, **r***^i^* is the response pattern in the i-th of *N_r_* runs, i.e. sets of measurements that are assumed to be independent from the other *N_r_* 1 sets. log of a vector here refers to the vectors of logarithms of each entry. * add a square root transform.

#### 2.0.1 Euclidean distances

The simplest distance to use is the Euclidean distance in the multivariate response space and this is an entirely valid choice. One can also use the squared Euclidean distance, which has the advantage that distances add over orthogonal dimensions. This additivity makes it particularly simple to model distances that arise from multiple underlying dimensions or parts of a model.

#### Correlation and mean-removed distances: normalizing population-mean activity and pattern variance

The population-mean dimension is the axis in the multivariate response space defined by the all-1 vector. Passing through the origin, this axis is at equal angles to all the axes corresponding to the measurement channels (e.g. neurons, voxels, electrophysiological sources). We refer to this axis as the population-mean dimension because computing the regional-mean by averaging activity across channels amounts to projection onto this axis (up to a scaling factor related to the number of responses). Regional-mean activation is usually studied using univariate analyses, for example in fMRI studies. To make the multivariate pattern analyses orthogonal and complementary to univariate regional-mean analyses, we may choose to remove the mean from each pattern estimate. In fMRI, an additional motivation for removing the regional mean is that each voxel averages many neurons’ response, and so the neural population mean dimension is overrepresented in the fMRI activity pattern estimates [66]. Instead of computing Euclidean distances on the activity patterns, we may therefore compute Euclidean distances after removing the mean from each pattern. Of course we may be losing distinctions between experimental conditions if we remove the mean and this needs to be kept in mind when interpreting the results.

A popular way to measure representational distance is the Pearson correlation distance (method=‘correlation’), which is defined as 1 − *r*, where *r* is the Pearson correlation between the two pattern estimates across the measurement channels. The Pearson correlation distance equals half the squared Euclidean distance measured after normalizing each activity pattern. The normalization of each pattern involves removal of the mean and scaling to unit variance. While removing the mean may be well motivated (as explained in the previous paragraph), scaling the pattern to unit variance can cause confusion. Two patterns of small activity values, which are close in terms of Euclidean distance, can appear quite distinct [65]. For example, patterns that contain only noise because they are elicited by stimuli that do not drive any significant response will have *r* ≈ 0, and so the correlation distance will be large: 1 − *r* ≈ 1. The patterns, thus, have a high correlation distance, although the corresponding conditions cannot be decoded from them. This caveat needs to be kept in mind when using the correlation distance.

#### Mahalanobis and symmetrized-KL Poisson distance: measuring distances relative to the noise

If we are interested in the discriminability of the representational patterns to downstream decoders, then we should measure the distance relative to the noise in the activity patterns. Figure 3a shows a case in which two measurement channels are negatively correlated. If two conditions (blue and green) differ along this direction, then the distance should be smaller, to reflect the fact that a downstream decoder will have lower accuracy.

**Fig. 3.**
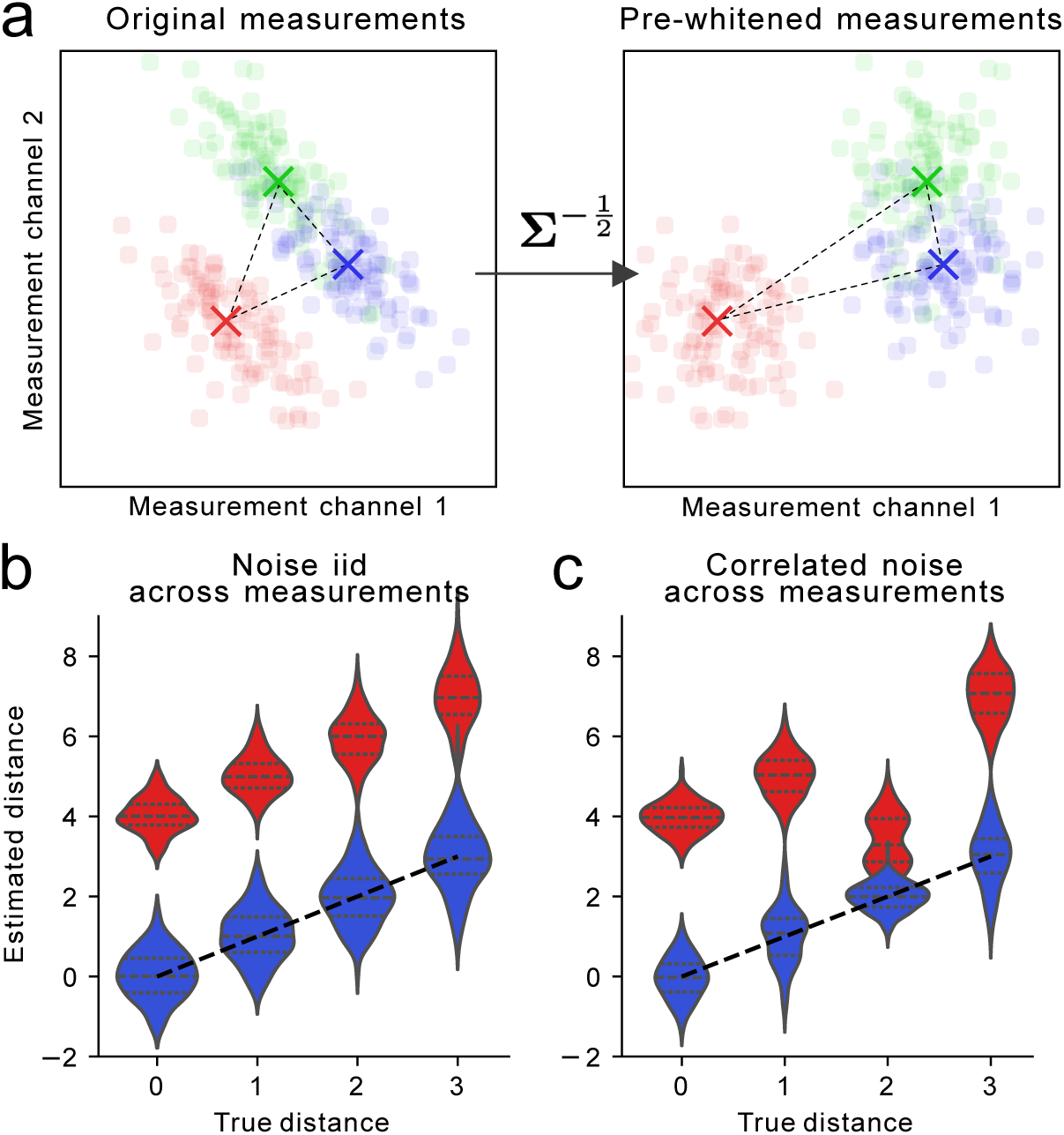
Impact of correlated noise across channels (a) and measurements (b,c) on distance estimation. **A**. Mean activity patterns for 3 conditions (red, green blue) plotted in the space of 2 measurement channels. In the raw data space (left), the two channels are negatively correlated across individual measurements (dots). After spatial pre-whitening (arrow), the correlation is removed. The distances in prewhitened space now reflect discriminability of the different conditions across measurements. **B**. Normal (red) and cross-validated (blue) estimation of four squared Euclidean distances (x-axis). When the noise is independently and identically distributed (iid) across measurements, the bias is constant for all squared distances and the rank-ordering of the distances is preserved. **C** Same as B, but the pair of condition with distance 2 has correlated (*r* = 0.5) measurement noise. This induces a negative bias in the distance estimate, also changing the rank-ordering of the distances. Cross-validated distances (blue) remove this bias.

If the noise distribution is multivariate normal, we can use an inverse square-root of the noise covariance matrix to transform the patterns into a whitened space, in which the noise is isotropic across measurement channels (Figure 3a). The distances in this new space are the Mahalanobis distances (method=‘mahalanobis’) between the original patterns. This procedure is especially adequate for fMRI data, for which the measurement noise is well approximated by a multivariate normal distribution, and for which there is often considerable correlation between measurement channels (voxels). For such data, multivariate pre-whitening has been shown to increase the reliability of the distance estimators [65].

For other measurement modalities, the noise may be non-normal and/or heteroscedastic. In this case, a variance-stabilizing transform can be used before measuring distances [15, 59]. For example, if we have neural data and assume Poisson noise, we can use a square-root transform on the spike rates to stabilize the variance. An important general approach to measuring the distance in terms of condition discriminability is the symmetrized Kullback-Leibler (KL) divergence. Each condition is represented by the distribution of patterns (reflecting the structured noise) in the multivariate activity space. The symmetrized KL divergence reflects the degree to which a response pattern drawn from either condition is decodable (in terms of expected log odds). The RSA3 Toolbox implements the symmetrized-KL dissimilarity estimator for Poisson noise (method=‘poisson’), which is applicable to patterns of neural firing rates. This is a close analogue of using the squared Mahalanobis distance for fMRI data, which is the symmetrized KL divergence between two Gaussian distributions with equal covariance matrix.

#### Removing the positive bias of distance estimates

When applied to noisy activity estimates, all distance estimators considered so far are positively biased in the sense that they tend to overestimate distances between the true noise-free activity patterns. Measurement noise perturbs response patterns in random directions. In high-dimensional space most of the noise is orthogonal to the line connecting the two patterns and thus tends to increases the distance between them.

If the measurement noise is independent and identical across conditions, the bias for squared distances is the same for all pairs of conditions (Figure 3b). However, if the noise variance varies across conditions, or if measurements for different conditions are correlated, the bias can be uneven across distances, such that even the rank ordering of the distances is affected (Figure 3c).

The bias can be removed by using crossvalidated distance estimates if repeated independent measurements of each pattern are available [65, 112–114]. These distance estimators multiply differences from different independent partitions of the data, which cancels the positive bias on average. This statement implies that if the true distance is zero, the mean cross-validated estimate is also zero (Figure 4b).

**Fig. 4.**
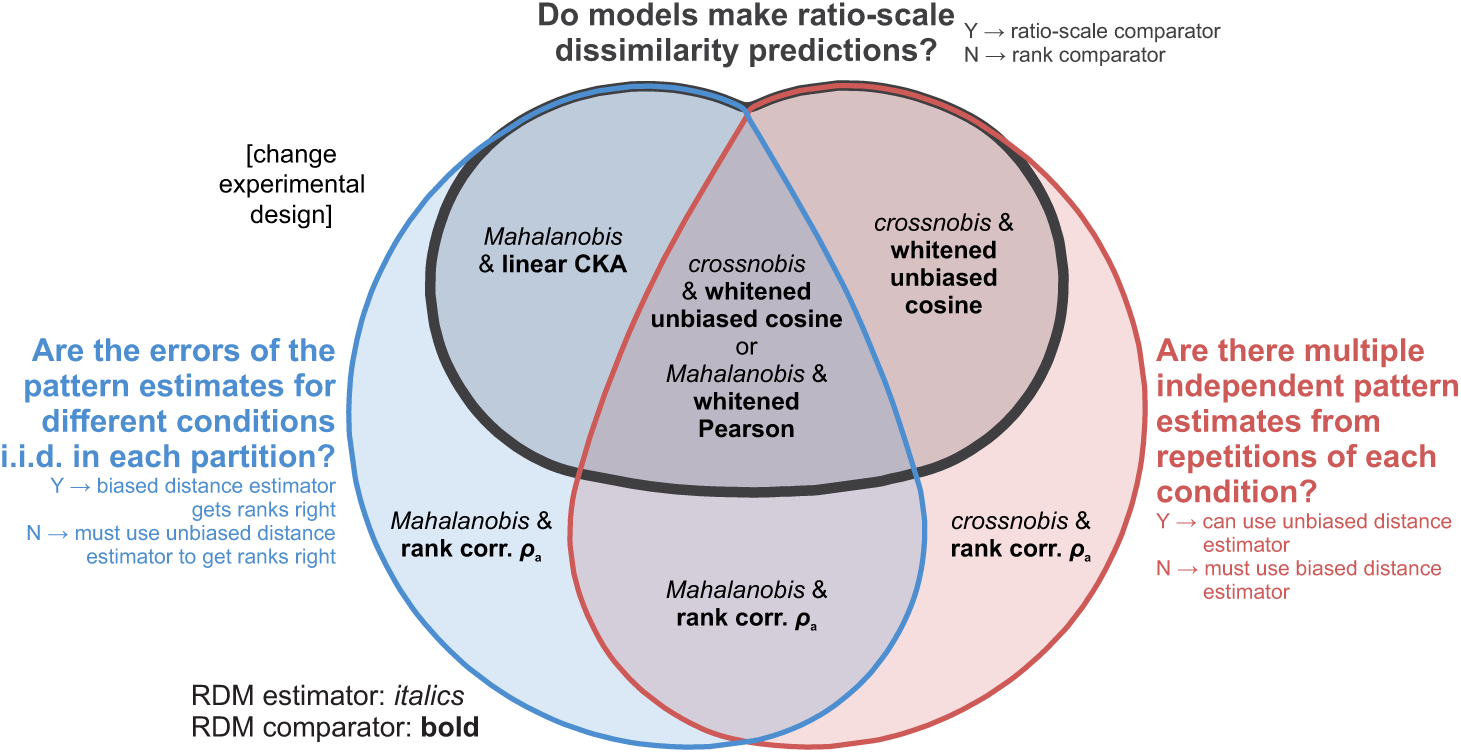
Choosing an appropriate dissimilarity estimator and RDM comparator. This Euler diagram shows which combination of RDM estimator and RDM comparator promises the most powerful model-comparative inferential analyses in different scenarios, and when a different experimental design is needed. To find the right combination of RDM estimator (italics) and RDM comparator (bold), answer the three questions (black, blue, red). For each question, a yes (Y) indicates that the answer is inside the set, and a no (N) indicates that the answer is outside the set. See text for details. The diagram is the minimum-contour-length iso-Euler diagram.

To achieve unbiasedness, the cross-validated estimator must sometimes result in negative values to average out positive estimation errors. Because distances are by definition non-negative, we use the term representational *dissimilarity* instead as a more general concept encompassing all the estimators we might want to use to characterize the representational geometry. Importantly, negative dissimilarities should not be excluded or replaced by zeros, but rather included in the subsequent analyses to ensure unbiased inference (see below). The only exception to this rule are analyses that require a valid distance matrix to function, such as visualization using metric multidimensional scaling.

The RSA3 Toolbox implements crossvalidated estimators for a variety of distance measures, including the squared Eucidean and squared Mahalanobis (crossnobis) distance and the symmetrized KL-divergence for poisson distributions. The crossnobis estimator (method=‘crossnobis’) is closely related to linear decoding analyses using the Fisher linear discriminant. In essence, it measures the distance after projection of test data points onto the Fisher linear discriminant dimension estimated with a set of training data points. Similarly, the symmetrized KL-divergence can be crossvalidated by using a different part of the data to estimate the probabilities used for weighting than for estimating the log-likelihood ratio (method=‘poisson_cv’).

#### Transformations on RDMs

Some research suggests that applying a non-linear transform to the distances can be advantageous. This enables the use of unsquared euclidean distances, which have stronger relationships to statistical measures of independence like the Hilbert-Schmidt independence criterion [115] and distance correlations [116]. Also a geo-topological transform can enable a stronger focus on the topological structure of the representation rather than its geometry [69].

We implement this with a general transformation function that allows the application of any transformation to the RDM and a specialized function for taking the square root and applying the geo-topological transform. One should apply analogous transformations to any pair of RDMs that one wants to compare.

### Step 3: Comparing RDMs

To draw these conclusions, we first need to choose a measure for the similarity of RDMs. Then we need to estimate how variable these similarity values are and how well a model could perform in principle. Once we have those estimates we can perform frequentist statistical inferences that test whether a model could be true, predicts anything about the data RDM, and for each other model, whether it significantly outperforms this other model.

#### RDM comparators

To compare RDMs to each other, in principle any measure of similarity for symmetric matrices or vectors can be used. However, different *RDM comparators* will weigh different aspects of RDMs differently. Most RDM comparators also have some aspects of RDMs that they are entirely invariant to, which is desired when those aspects are deemed irrelevant for the research question. Beyond that, a different weighting of the RDM dimensions will still lead to different evaluations - that is, different RDM comparators maybe more sensitive to measurement noise or differences between subjects. We give an overview of the possible RDM comparators in the toolbox in Table 2.

**Table 2.**
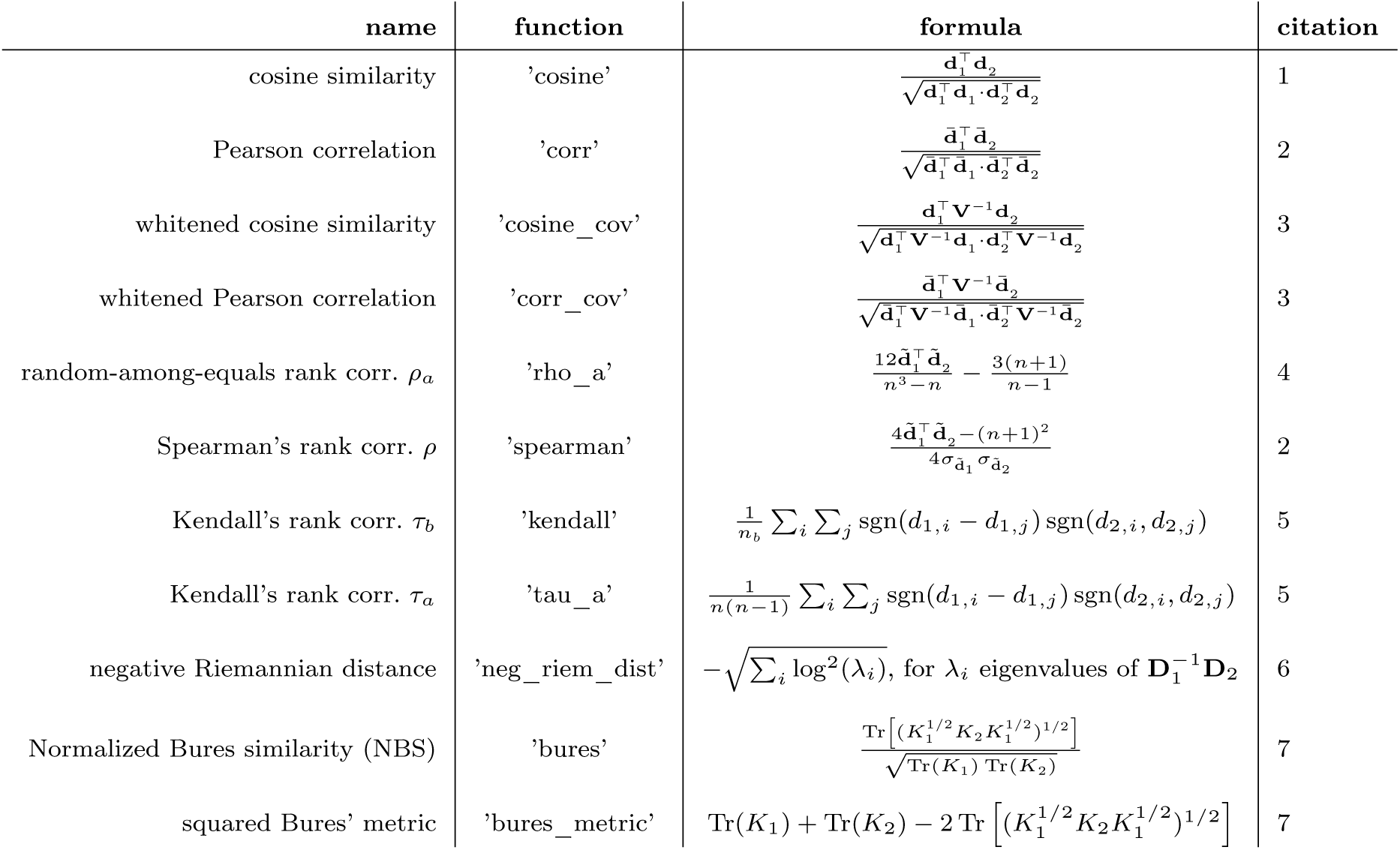
Methods for comparing two RDMs to evaluate predictions of representational geometries. The column labeled “function” specifies the name to be passed to the compare function to compute the similarity. In the formulae, **d***_k_* for *k* = 1, 2 are the two vectorized RDMs, *d_k,i_* is the *i*-th scalar dissimilarity of **d***_k_*,**d̄**^^*k* is the centered version of **d***_k_* (with the mean subtracted from each dissimilarity), **d**°*k* is the rank transformed version of **d***_k_* and **D***_k_* are their transforms into positive definite matrices as described in the text. *n* is the number of dissimilarities, *n_b_* = *^√^n_b_*_1_*n_b_*_2_ is the normalization for *τ_b_* where *n_b_*_1_ and *n_b_*_2_ are the numbers of ordered pairs in the two compared distance vectors. **V** is the *n × n* covariance matrix of the dissimilarity estimate errors. Citations: 1. [114] 2. [17] 3. [70] 4. [71] 5. [113] 6. [67] 7. [117] .

**Table 3.**
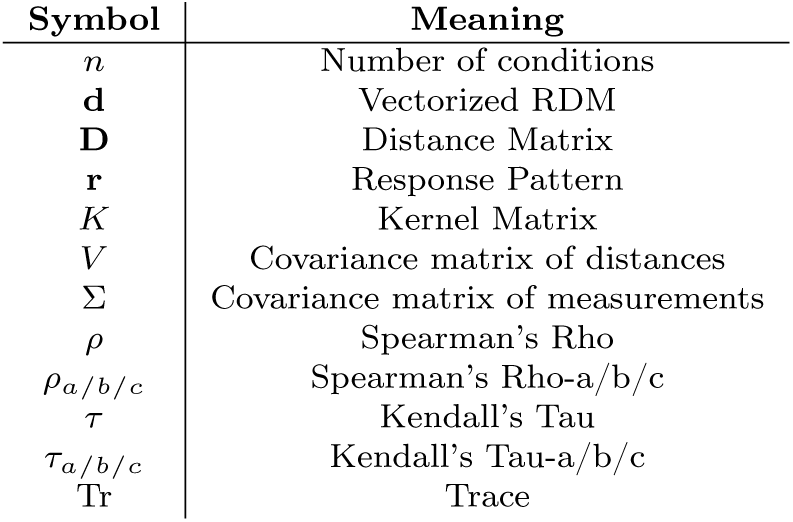
Symbols used.

| Symbol | Meaning |
| --- | --- |
| $n$ | Number of conditions |
| $\mathbf{d}$ | Vectorized RDM |
| $\mathbf{D}$ | Distance Matrix |
| $\mathbf{r}$ | Response Pattern |
| $K$ | Kernel Matrix |
| $V$ | Covariance matrix of distances |
| $\Sigma$ | Covariance matrix of measurements |
| $\rho$ | Spearman's Rho |
| $\rho_{a/b/c}$ | Spearman's Rho-a/b/c |
| $\tau$ | Kendall's Tau |
| $\tau_{a/b/c}$ | Kendall's Tau-a/b/c |
| Tr | Trace |

Generally, we want to ignore the overall scale of the RDM to make our inference independent of the overall signal strength, which often varies across subjects, sessions, and measurement modalities. Thus, we want all our RDM comparators to be invariant to an overall rescaling of the RDM. Formally, *sim*(**d**_1_, **d**_2_) = *sim*(*a***d**_1_*, b***d**_2_), for all *a, b* ∈ ℝ^+^.

##### Cosine Similarity

The simplest measure of similarity that ignores the scale is the *cosine similarity* of the vectorized RDMs. This similarity is the inner product of the vectorized RDMs after dividing each by their norm. This normalization explicitly removes the dependence on the scale of the RDMs. However, the cosine similarity is sensitive to the average distance value, i.e. adding a constant to all distances changes cosine similarities. In particular, the bias introduced by estimating distances from noisy data reduces the cosine similarity below 1, even if the bias is constant and all representations become more similar if a constant is added to all distances in all representations. For our example in Fig. 3b, the cosine similarity of the expected RDM estimate is XXX. Thus, the cosine similarity should only be used for cross-validated or noise free dissimilarity estimates.

##### Pearson Correlation

To remove the assumption that a model-predicted distance of 0 corresponds to a measured dissimilarity of 0, we can use the Pearson correlation coefficient *r* between RDMs. Like the cosine similarity, the correlation is the inner product of the vectorized RDMs after normalization. But the normalization includes mean subtraction as well as division by the norm. It is thus equivalent to the cosine similarity after subtracting the average dissimilarity.

##### Whitened RDM similarities

Cosine similarity and Pearson correlation are adequate measures of similarity of two vectors when the individual elements of the vectors are independent. The entries of a RDM are not independent, because all dissimilarities involving the same condition are based on the same measurements of that condition. For squared Euclidean, Mahalanobis, and Crossnobis dissimilarities, we can derive the covariance matrix *V* of all the dissimilarities analytically [70]. This estimate can then be used to “whiten” - i.e. to de-correlate - the dimensions of the RDM (see Table 2). We have shown that model comparison using these whitened versions of the cosine similarity of Pearson correlations outperforms model comparison using traditional methods [70].

Interestingly, the whitened cosine similarity is equivalent to the linear centered kernel alignment (CKA) [63], if the measurement noise within each condition is assumed to be iid. This equivalent formulation can be computed faster as it avoids the inversion of *V* . Our implementation uses this equivalent formulation for faster computation.

##### Rank correlation coefficients

We can drop the assumption of a linear relationship between RDMs by using rank correlation coefficients like Kendall’s *τ* or Spearman’s *ρ*. For this lowest bar for a relationship, Kendall’s *τ_a_* or *ρ_a_* are preferred over the standard Spearman’s *ρ* or Kendall’s *τ_b_*and *τ_c_* because the latter all favor models that predict RDMs with tied ranks. The recommended option is *ρ_a_*, which is as computationally efficient as Spearman’s *ρ* and like Kendall’s *τ_a_* correctly handles models that predict tied dissimilarities. The rank correlation coefficient *ρ_a_* is the expectation of Spearman’s *ρ* under random tie breaking (using a closed-form solution).

Rank correlations are helpful if our models only predict the order of distances not a particular distance. For example, just that distances within a category should be smaller than the ones between categories.

##### Bures similarity and distance

Two alternative comparators for positive definite matrices are the Bures similarity and distance [117]. These metrics are equivalent to generalized shape metrics [62], which allow an orthogonal rotation to align the shapes with each other before measuring their distance. As these measures are based on the centered kernel matrices instead of distance matrices, we need to convert the distance matrices to kernel matrices first. Fortunately, distance matrices and centered kernel matrices contain equivalent information and the transformation between them is a simple linear map, such that this is not an issue.

##### Riemannian manifold similarity

Yet another measure of similarity for positive definite matrices is known as the Riemannian manifold distance [67]. This distance ensures that applying the same invertible linear map to the two activity matrices does not change their distance. Additionally, the original publication showed promisingly high reliability of this measure. Main drawbacks are slight stability issues for close to singular distance matrices and a relatively long computation time.

##### Toolbox implementation

All comparison methods are implemented in rsatoolbox.rdm. They can each be accessed by passing a method argument to rsatoolbox.rdm.compare or by using a specific function rsatoolbox.rdm.compare_[comparison]. The comparison functions each take two RDMs objects as input and return a matrix of all pairwise comparisons.

#### Choosing an appropriate combination of RDM estimator and RDM comparator

We should choose the combination of RDM estimator and RDM comparator that promises the greatest power for inferential model comparisons. The best choice depends on the answers to three questions (Fig. 4):

**(1) Do the models make ratio-scale dissimilarity predictions?** If yes (black set), we can gain power by using a ratio-scale RDM comparator. Such a comparator measures to what extent the measured dissimilarities are proportional to the model-predicted dissimilarities, taking 0 dissimilarity predictions to mean that the measured dissimilarity should also be 0. If the answer no (the model does not make ratio-scale predictions; outside the black set), we can evaluate the model on the basis of the ranks of its dissimilarity predictions, using a rank correlation coefficient. The *ρ_a_* coefficient is an appropriate and computationally efficient choice.
**(2) Are the errors of the pattern estimates for different conditions independent and identically distributed (i.i.d.) within each partition?** If yes (blue set), then a biased distance estimator (without crossvalidation) will get the ranks right, as the noise-induced bias will be constant across squared distances (Fig 3b). If no (outside blue set), different levels of error variance between conditions and dependency in the error variance between conditions can compromise the dissimilarity ranks when using a biased RDM estimator (Fig 3c), so an unbiased (crossvalidated) RDM estimator is needed.
**(3) Are there multiple independent pattern estimates from repetitions of each condition?** If yes (red set), it is possible to use a crossvalidated RDM estimator and thus obtain unbiased distance estimates. If no (outside red set), we do not have the repeated measurements needed for crossvalidation and can only obtain biased distance estimates.

In case the answers to questions (2) and (3) are both no (outside both red and blue sets), there is no way to correctly estimate even the ranks of the dissimilarities, so it is best to use a different experimental design. In the central intersection of all three conditions, both approaches work with different advantages: The combination of the crossnobis estimator and the whitened unbiased cosine similarity comparator promises an interpretable 0 point and no bias, whereas the combination of the Mahalanobis distance and whitened Pearson correlation comparator promises slightly lower variance [70]. Which of these two choices affords more power in model adjudication depends on the model predicted RDMs (if the placement of the 0 point on the distance scale differs between models this fact favors the crossnobis estimator) and the proportion of data that must be held out in crossvalidation for the crossnobis estimator (more favors the biased estimators)

### Step 4: Inferential model comparisons

Our inference about our models usually aims for two things: First, we want to compare each model to the data and judge how well the model captures the representational geometry we observe in the data. In particular, does it capture any aspect of the representational structure at all? How good is the fit given the level of measurement noise? Second, we want to compare models, i.e., draw conclusions about whether the difference between two models could have happened due to chance even if the two models were in truth equally good.

#### Model specification: Selecting a model type and defining the RDM model

To evaluate hypotheses we need to capture those hypotheses into models. In the rsatoolbox we conceptualize models as objects with two main functions: First, they can predict an RDM for the set of conditions we want to analyze, potentially based on parameters. Second, they provide a fitter method to find the best parameters for a given dataset. This setup is very broad, encompassing all parametric models of representational similarity.

The prediction function is implemented as functions of the model object in two flavors, predict_rdm returns the RDM object including descriptors. predict directly returns a ’naked’ RDM vector, which allows for slightly faster evaluations, if the descriptor annotations are not needed.

There are different methods for defining a model depending on what kind of flexibility is needed for the model. The first choice is whether any flexibility is required.

If the hypotheses can be captured by *fixed models*, i.e. with models that have no free parameters and predict a single fixed RDM, this simplifies the inference substantially as cross-validation is not needed then.

However, there are situations that require flexible models. In particular, many models contain some flexibility as to how their activity patterns are distorted by the measurement process, i.e. a flexible measurement model. A common example of this is the size of voxel averaging regions, which is typically unknown and distorts the RDM [41]. In such cases, we require a proper flexible model whose parameters are fit to the data, because model performance may otherwise depend primarily on luck with the parameter assignment [41].

To enable this kind of fitting, the rsatoolbox provides two broad approaches:

First, *selection* and *interpolation* models can be used for arbitrary relationships between parameters and RDMs, as long as the range of RDMs can be represented by a small selection of RDMs or by a one dimensional manifold of RDMs. These models work well for these cases and fitting is extremely stable. Thus, these models are recommended for situations similar to the fitting of voxel sizes.

The other approach is a weighted model, which aims to capture the weighting of feature dimensions or model components. The central idea here is that squared euclidean distances are additive for orthogonal axes, i.e. two conditions with squared distance 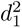 along a first dimension and 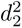 along a second dimension will have total squared distance 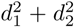. This connection allows us to combine the distances from different parts of a model into an overall distance. When we do not know how the two distances are weighted relative to each other, we can add a weight to each distance. The overall squared distance RDM is then simply the weighted sum of the component RDMs.

##### Fixed models

are models that predict a single RDM without any parameter. These models are the simplest and most frequently used type in RSA. To formally fit the model object architecture, these models have a dummy_fitter function that are always None and ignore any optional parameter to their prediction functions. To homogenize the analysis we generally expect these fixed model objects, not RDM objects as input to the inference methods described below.

##### Selection models

are models that are given a set of RDMs and predict that the true RDM is one of these RDMs. These models give a coarse simple method to implement an arbitrarily complex set of potential RDMs. Their prediction takes the index of the selected RDM as an input and they allow only one particular fitting function, which evaluates each RDM from the set on the training data and simply returns the index of the best performing RDM.

##### Interpolation models

interpret a given set of RDMs as the interpolation points for a piece-wise linear manifold of RDMs. This allows an approximate model for any one dimensional manifold of RDMs, i.e. any set of RDMs that is created by changing a single parameter. These models can be fit efficiently with another specialized fitting function, that first finds the best performing RDM in the set and then uses a bisection method for one dimensional optimization to find the best RDMs in the linear segments next to the start RDM. Bisection for one dimensional optimization converges quickly even without any gradient information.

##### Linear weighted models

predict the RDM as a weighted sum of a set of basis RDMs. This type of combination primarily appears for feature reweighting models, which assume that there are underlying (groups of) features, whose relative contributions to the representational dissimilarities are unknown to the experimenter. If features are independently scaled by *w_i_* the resulting euclidean RDM is the sum of feature RDMs weighted by *w*^2^. Based on this observation, we usually want to impose a positivity constraint on the weights as there is no possible feature weighting that would push conditions together that are separated along a particular feature. Linear models require a separate real parameter for each basis RDM. Thus, they often require regularization to be applied successfully. To accommodate these different fitting and regularization methods we provide different fitting functions. In particular we provide linear algebra based methods that fit the RDM based on euclidean, cosine or correlation similarities (described below) and optimization based methods to include the positivity constraint and *L*_1_ and elastic net regularization.

##### Arbitrary custom fitted models

These model types capture only a small selection of possible RDM models although they already expand the model zoo substantially beyond what earlier toolboxes provided. If users are keen to implement other model types our toolbox can be expanded easily. As long as the new model object implements the prediction and fitting methods, it can be used in all inference methods we provide in the toolbox.

#### Model identification and evaluation

All flexible model types (i.e. all but the fixed model type) have parameters that need to be fitted to data. Fitting a model’s parameters to the data is known as model identification because it identifies the particular instance of the model (i.e. the settings of the parameters) that fit best. Because a fitted model is always somewhat overfitted to the fitting data, the goodness of fit on the data used for fitting is not an unbiased measure of model performance. Model evaluation therefore requires a test of the fitted model on data not used in model fitting. To make good use of limited data, we perform model fitting and performance evaluation in crossvalidation. This approach yields good estimates of the performance each fitted model achieves on new data, i.e. of the model’s generalization performance. The resulting performance estimates can also be compared between models that have different degrees of flexibility (e.g. a fixed model, a low-parametric model, and a high-parametric model).

We need to decide what type of generalization performance to require. If all our models are fixed, there is no fitting, and thus no overfitting to account for and cross-validation is not needed at all. In a case study of the tactile representations of the five digits of the right hand in the brain of a patient, we might fit and evaluate the models on separate brain-activity data sets in which each of the five fingers has been stimulated (same subject, same conditions). In a group study of sensory representations of digits, we might want our models to generalize to new subjects. We would then evaluate each model on data from different subjects acquired during stimulation of the same five fingers (different subjects, same conditions). In a case study of visual representations of face images in a single prosopagnosia patient, we might want our models to generalize to new face images (just like computer vision models must work on new stimuli) to test to what extent they can account for the representations underlying face perception in the patient (same subject, different conditions). Finally, in a group study of 24 people’s visual representations of object images, we might want our models to generalize to new subjects and new images simultaneously (different subjects, different conditions). The RSA3 toolbox supports model evaluation for all these types of generalization. The type of generalization should always be reported and carefully considered in interpreting the results.

##### Cross-validation

Once we use flexible models, we need to worry about overfitting, as more flexible models will fit data used for fitting their parameters better, even if that flexibility does not correspond to any real effect. This is a well known problem for statistical model comparisons of any kind and many correction methods for this problem exist. For RSA, the most common technique to solve this problem is cross-validation, where we split the data into groups, which are in turn used for evaluation, while all other data are used as a training set to fit the model parameters. This yields an estimate of model performance that does not overly favor flexible models anymore.

For RSA the overfitting argument applies to both subjects and stimuli. For cross-validation the question is how far the model should generalize. For example, if we use some stimuli for fitting and others for evaluation, good model performance requires the model to generalize to new stimuli. This is a separate question from the generalization we aim for with our bootstrapping procedures. By bootstrapping we aimed for our inference to generalize to new subjects or stimuli. In some situations our models may require subject or stimulus specific parameters, but we can consistently find parameters for each separate subject or stimulus set that work. If we ask our models to generalize to new subjects or stimuli, we do most likely want our inferences to generalize to new samples along that factor, too though.

In our toolbox we implement cross-validation in two steps: We first split the dataset into the cross-validation folds (see inference.sets_*) and then apply the cross-validation function (inference.crossval). Furthermore, the bootstrapping functions do contain additional parameters to implement cross-validation within the bootstrap (inference.eval_bootstrap_*).

#### Model-comparative inference

To compare different models to each other, we need to estimate how uncertain our experimental results are. For our purposes, this uncertainty is quantified by a covariance matrix of the model evaluations over repetitions of the experiment. A key question for computing this covariance is how far we want our results to generalize, i.e. which aspects we assume to vary over the repetitions of the experiment. This generalization of our inference is different from the generalization we ask our models to perform and test with crossvalidation.

##### Inter-subject variability

The simplest type of generalization for RSA is to generalize to new subjects performing the same task with the same stimuli, because all evaluations we use are averages across subjects. The covariance of this average model performance then simply is <u>^1^</u> times the covariance of model evaluations over subjects for which we can use the standard sample estimates. In this situation, simple t-tests and rank-sum tests provide adequate tests to test whether two models perform significantly differently well. These tests are by far the most common type of inference used for RSA.

##### Two-factor bootstrap

In most cases we also want to generalize to new stimuli used for evaluation [71, 118]. As our model evaluations are not means across stimuli, we cannot easily compute the covariance of the model performances if we sampled new stimuli. Instead, we can resort to bootstrapping methods. If we want to generalize only to new stimuli, standard bootstrapping across conditions produces accurate estimates of the variance due to a random choice of conditions. Technically, bootstrapping subjects also reproduces the covariance due to the random sampling of subjects. There is no advantage of using the bootstrap over the direct formula for this variability though such that bootstrapping is not used for the variability due to subject choice in practice.

Once we want to generalize across both subjects and conditions, a naive bootstrap approach yields substantially too large variance estimates [71] due to roughly triple counting measurement noise and any other variance that is not due to the subject or stimulus choices. To counteract this effect, we can use a correction formula that results in fairly good variance estimates for this two factor bootstrap.

In the toolbox the bootstrap methods are available in the inference submodule as eval_bootstrap_* functions.

The bootstrap is compatible with crossvalidation to correct for overfitting of flexible models [71]. We refer to the combination as bootstrap wrapped crossvalidation. Due to the bootstrap resampling we usually cannot keep the crossvalidation folds constant across bootstrap samples, which requires another correction [71]. This functionality is available through the eval_dual_bootstrap function in the toolbox.

##### Statistical tests

Once we have estimated the model performances and our uncertainty about them, we can use this information for frequentist statistical tests that test whether two models perform differently well, whether a model performs above chance and whether a model performs worse than the noise ceiling. In our toolbox, we mostly recommend using t-tests based on the means and the covariance of model performances, which can be estimated for different generalizations as discussed above. In the toolbox, this functionality is implemented as functions that apply to the results objects that all evaluation functions yield as outputs as these tests do not require much computation. The toolbox also supports classic rank based tests. One should note that these tests are based exclusively on the variance due to subjects and cannot be used to take the variance due to condition choice into account.

#### Visualizing model comparisons

Visualization has an important role because it is needed to help researchers make sense of a complex set of results. Theoretical progress depends on comparing many models. The toolbox offers the functions plot_model_comparison and map_model_comparison, which visualize the quantitative and inferential results, revealing: (1) how well each model performs, (2) which other models each model significantly dominates, (3) which models explain significant variance, (4) which models leave significant explainable variance unexplained (do not approach the noise ceiling), (5) how similar the predictions made by the models are to each other, and (6) how similar activity patterns are to each other across multiple brain regions.

Fig. 5 shows simulation results visualized with the two visualization functions. All statistical tests are adjusted for multiple testing, controlling either the false-discovery rate or the familywise error rate. The **bar plot** (plot_model_comparison, Fig. 5a) integrates points (1) to (4). This provides an essential basic visualization integrating quantitative and inferential results. Key elements include the *noise ceiling* (gray bar), the “*dew drops*” (white and gray half circles indicating significant differences of model performance from 0 and from the lower bound of the noise ceiling, respectively), and the “*model-dominance wings*”, which show which other models each model dominates. The toolbox gives different options for visualizing the pairwise inferential comparisons. One option is “*non-significance cliques*”, which communicates all pairwise comparisons in terms of a minimal set of model cliques within which no differences are significant. Another option is “*double arrows*”, which communicates all pairwise comparisons by a minimal set of double arrows. Each of a set of horizontal double arrows indicates that all models to the left of it are significantly different from all models on the right of it. This approach is visually efficient (few elements) when the researcher chooses to plot the bars in ascending or descending order of model performance. These practical advances become important when a larger number of models is compared. For example, for just 40 models, there are 780 pairwise comparisons.

**Fig. 5.**
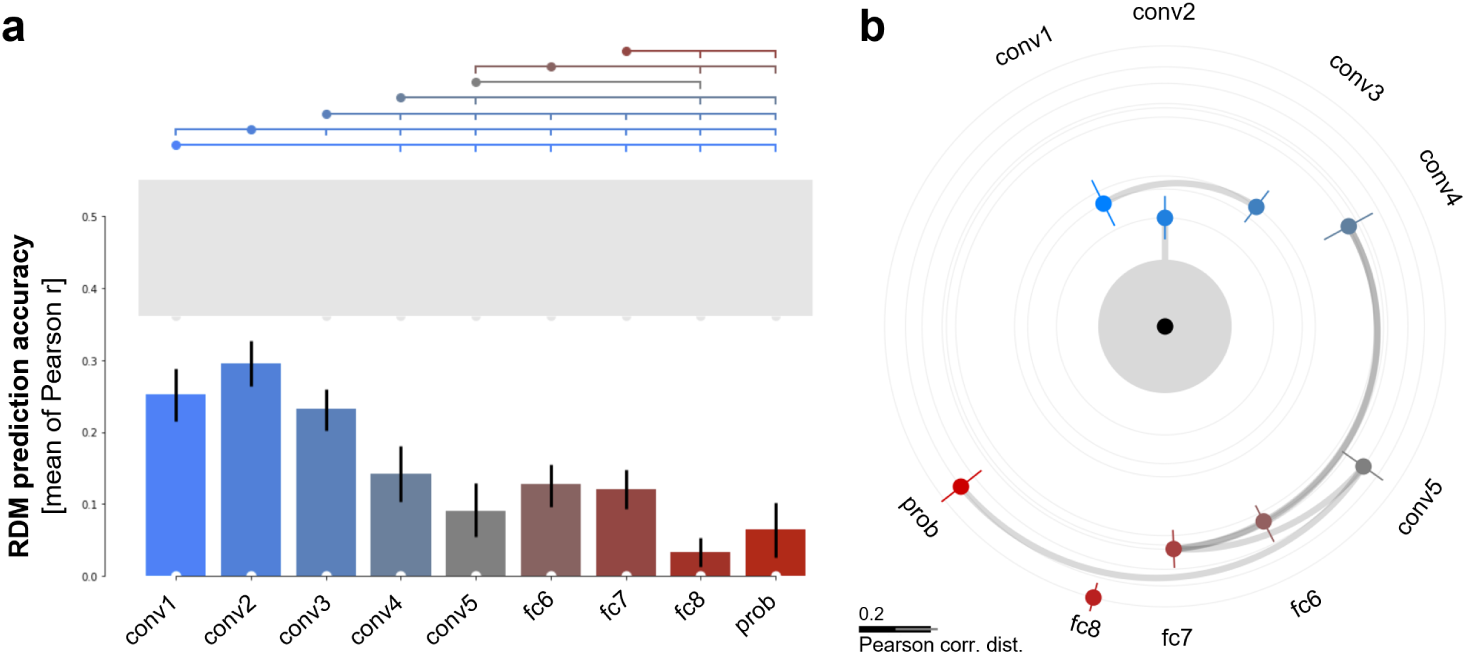
Visualization of model-comparative inference results. Results of inferential analysis of simulated data. Ground-truth model is convolutional layer 2 (conv2) of AlexNet. All layers of AlexNet (blue to red) serve as candidate models. **(a) Bar plot of RDM prediction accuracies.** Model comparisons two-tailed, false-discovery rate controlled at *q <* 0.01 (36 model-pair comparisons). Inference by bootstrap resampling (1,000 samples) of subjects. Error bars are 95% confidence intervals. One-sided comparisons of each model performance against 0 (white “dew drops” at the bottom indicate significant difference from 0) and against the lower-bound estimate of the noise ceiling (gray bar, gray dew drops indicate significant difference from the noise ceiling) are Bonferroni-corrected for 9 models. “Model-dominance wings” (top) indicate, for each model (dot in model color), which other models it significantly dominates (downward tick marks). **(b) Model map.** Same results as (**a**), but deviations of model predictions from the data are used to map the models around the data RDM with a modified multidimensional scaling (MDS). Inter-RDM distances measured by the Pearson correlation distance. MDS constrained to represent the deviations from the data RDM exactly (same information as in **a**) and the deviations among model RDMs (not shown in **a**) approximately.

**Fig. 6.**
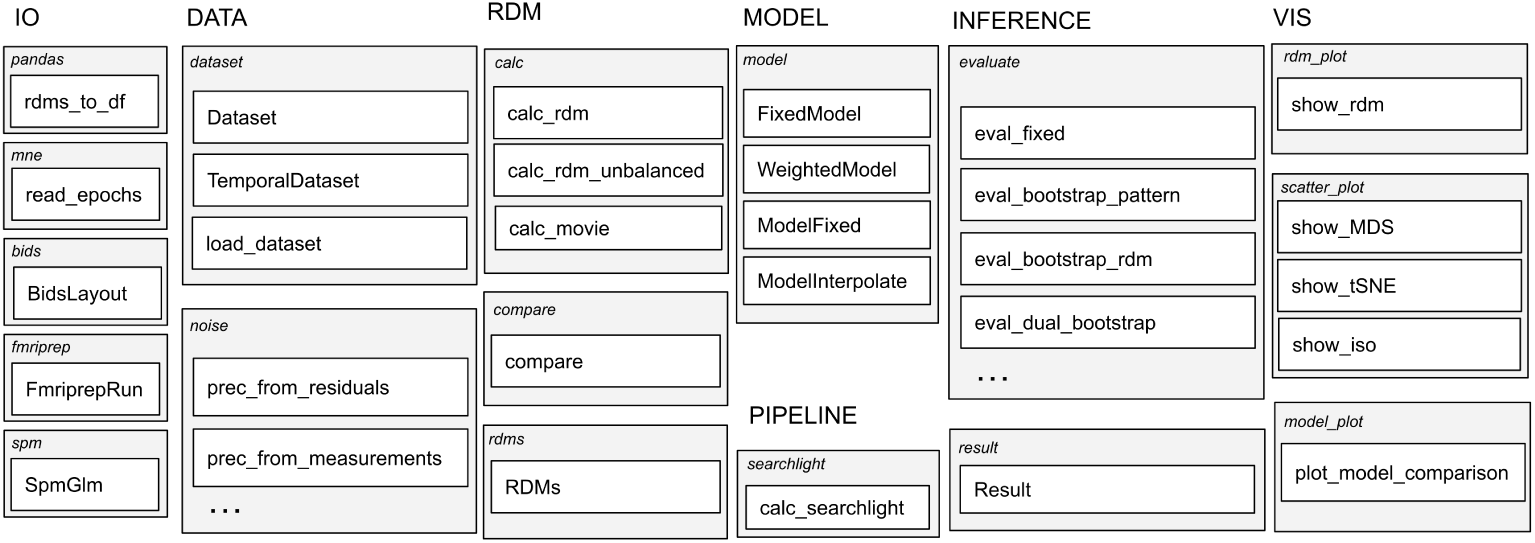
Library overview. Structure of the library with key elements listed. The columns display the sub packages of rsatoolbox; corresponding from left-to-right with the typical order in which they come into play during an analysis. Light gray grouping boxes represent modules annotated with the respective module name. White boxes are key functions (lowercase) and classes (capitalized nouns).

The **model map** (function map_model_comparison, Fig. 5b contains all the information in the bar plot, points (1)-(4) above, but also shows approximately how similar the predictions of different models are to each other (5). Distances in this map reflect deviations among the RDMs. The central black dot is the data RDM. The noise ceiling becomes a “*noise halo*”. The best model (conv2 here) is, by definition, shown straight above the data RDM. The analysis correctly identified the true model (conv2) in this simulated data set. Gray arcs connect models that are not significantly different. Conv2 is not statistically distinct from the noise halo (vertical gray arc). The arrangement is computed with multidimensional scaling (MDS) using the metric stress criterion. However, the deviations of the model RDMs from the data RDM (radii) are constrained to be exactly represented. The scale bar shows the length corresponding to 0.2 Pearson correlation distance units, and its error bar shows the full range of %-errors incurred by mapping into 2D. We will refine this visualization by adding a radial scale (to replace the scale bar) and different options for the cost function that the mapping minimizes.

### Step 5: Multiple testing across space and time

In some scenarios we do not have a specific hypothesis about where in the brain or when in time experimental effects may occur. This requires a more exploratory approach: instead of applying the above analysis steps to data from a particular brain region or time window of interest, we may opt to systematically scan brain space or time in search for effects. This exploratory approach involves repeated inference across spatial locations or time points. Below, we describe how the toolbox facilitates such analyses.

#### Searchlight inferential mapping

In the spatial domain, a popular approach uses a “searchlight” to scan the brain for experimental effects [119, 120]. A searchlight is a small analysis volume that covers a local neighborhood in brain space. The searchlight is systematically moved across the brain, centering it on each location in turn. This process repeats the analysis at each location and yields an inferential map showing where in the brain representational effects occur. This method promotes the discovery of brain regions that encode information about the experimental conditions (e.g., [112]) or whose representational content can be explained by a conceptual or computational model (e.g., [121, 122]). Searchlight inferential mapping can also be applied on the cortical surface for better spatial selectivity and increased sensitivity to local information content [123]. The toolbox supports both volume and surface-based searchlight mapping.

#### Representational dynamics

In the temporal domain, it is common to use a sliding window approach to examine representational dynamics [38, 42, 124–126]. In this approach, the researcher defines a temporal search window and slides the window along the time axis of the data. For each time window, spatial response patterns are extracted from sensor or source space. This process repeats the analysis at each time point and yields an inferential time course showing when in time representational effects occur. This method, sometimes referred to as a (spatio)temporal searchlight or sliding window analysis, is useful for investigating dynamic changes in representational content, such as when representational distinctions emerge [38, 124, 127] or when a model best aligns with neural data [25, 55, 125, 128].

The rsatoolbox does not contain internal methods to handle correct inference with many correlated results over space or time. There are well established methods for this kind of multiple comparisons problem (MCP) as implemented for example in Nipype [75] or MNEpython [76]. For analyses of representational geometries over time, we recommend applying such external MCP correction tools on the uncorrected similarity results obtained from our toolbox.

## 3 Discussion

This paper presents a new toolbox that implements RSA in Python, along with a third wave of methodological progress [70, 71]. The toolbox consolidates previous developments and brings the following advances:

1. **Whitened RDM comparators.** Whitened RDM comparators (whitened versions of the cosine and Pearson RDM correlation for biased and unbiased dissimilarity estimators) generalize the linear centered kernel alignment [63] to unbiased distance estimators, approximate a likelihood-based criterion for model selection, and provide near-optimal sensitivity to subtle differences between models [70].
2. **Model-comparative inference generalizing to the populations of subjects and conditions.** Fixed or parameterized models can now be compared with an inference procedure that generalizes to populations of subjects and/or conditions [71, 129]. The key innovation is a new 2-factor bootstrap procedure for inference on parameterized models [71]. Models are fitted and evaluated in crossvalidation avoiding bias caused by overfitting to either subjects or conditions, while the inference can treat subject, condition, or both as random effects. The new procedure is substantially more powerful than the earlier double bootstrap [113, 129] and has been validated with simulated and real data (Calcium imaging and fMRI) [71]. The new inference framework supports the novel whitened RDM comparators as well as a range of other RDM comparators, including rank-correlation coefficients, which are less sensitive, but have the advantage that they require weaker assumptions for valid inference.
3. **Dissimilarity estimators for neural data.** We introduce several dissimilarity estimators specifically designed for neural data (based on the Poisson distribution). In particular, the Poisson symmetrized-KL-divergence estimator and its crossvalidated variant provide improved RDM estimates on the basis of neural recording data.
4. **Efficient rank-based model evaluation.** We introduce Kendall’s *ρ_a_* [130], a rarely used Spearman-type alternative to Kendall’s *τ_a_*, which like the latter correctly treats models that predict ties [113], but is much faster to compute. The efficiency of *ρ_a_* is essential when we are comparing many models or mapping inferential results across the brain with searchlight RSA.
5. **Model refutation by comparison to the lower bound of the noise ceiling.** Inferential model comparisons to the lower bound of the noise ceiling are now available
6. **Better visualization of model comparisons.** To more concisely present inferential results of all pairwise model comparisons, novel alternatives to “Nili bars” [113] include “Golan wings” and “double arrows” (Fig. 5) or “cliques” of models whose performance does not differ significantly. Furthermore comparisons to the noise ceiling and to 0 or chance-level performance are visualized using “dew drops” or “icicles”. The new options can more efficiently summarize all model-comparative inferential results, which becomes essential as the number of models grows.
7. **Model map of performance, inferential comparisons, and model relationships.** A novel “model map” (Fig. 5) visualizes the descriptive and inferential model comparisons in a 2D diagram, with the data RDM at the center within a noise halo (in place of the noise ceiling). Radial distance of each model exactly represents model prediction error, and the polar angles of the models approximately reflect the similarities among the models’ predicted RDMs. Connections between points link models whose performance does not differ significantly, defining all pairwise model-comparative inferential results.

### 3.1 Future plans

We will continue to maintain and further develop the rsatoolbox and invite others to include any coming developments into the toolbox.

There are a number of features that could be added to further improve convenience of use. In line with other analysis tools in the neuroscience ecosystem, we plan to encapsulate the toolbox into a RSAtoolbox BIDS app, and to provide a fully functional container image with a simple interface that can be slotted into existing data processing pipelines. In order to further support the reproducibility of projects based on rsatoolbox, such containers or apps will annotate the steps taken in a standard format [131, 132]. Encapsulation will also require support for a larger variety of data formats for import.

Additionally, better visualizations will always remain a welcome addition. We will extend the model map into a 3D interactive visualization. We will develop a visualization that summarizes the relationships among multiple data RDMs estimates from neural populations in different brain regions and multiple model RDMs in a 2D or 3D map. Such a visualization will be extremely useful for giving the researcher a sense of the transformations of representational geometries across stages of representation in brains and models and about the global relationships between model and brain representations.

Finally, we expect that new additions and variations to RSA will still be developed. Whenever that happens, we aim to extend the toolbox to include such new additions. Such new developments could include: New methods for constructing RDMs, new methods for comparing RDMs, new methods to perform statistical inference, new APIs to handle different data inputs or outputs, and many more.

### 3.2 Software development philosophy

To enable future additions and extensions to the toolbox and integrate it well with other tools in neuroscience, we converged on a number of principles for the development of the rsatoolbox:

We build our toolbox to include all commonly needed functionalities for RSA analyses, going from the preprocessed data to the final inference. To enable extensions, we aim to make the code as modular and reusable as possible. This will allow users to easily replace elements of their pipeline, and contribute new developments to the toolbox. As a standard for inclusion, we require all core statistical methods implemented in releases of the toolbox to be described and evaluated in peer-reviewed publications. More experimental approaches can be implemented in non-main branches, such that this criterion does not impede the speed of development, while providing a clear and transparent quality standard for the main release.

To enable community-driven development, we follow best practices in version control (Github), code review, continuous Integration [133], packaging (PyPi), and documentation (readthedocs). Users can submit queries, feature requests and bug reports by raising Issues on Github, which are closely tracked. These measures ensure accurate and reproducible analysis and enables discussions to reach a consensus about the best implementation for a given feature. Additionally, we document the current best practices in our documentation: Auto-generated API documentation is generated from in-code docstrings and key functionality is accompanied by narrative documentation as well as full demos in Jupyter Notebooks. The full documentation is then build and hosted at readthedocs (https://rsatoolbox.readthedocs.io/). Such complete, accessible documentation is key to the ease of use of software.

The library is shared under the MIT License (https://opensource.org/licenses/ MIT), because of its simplicity, permissiveness and compatibility with other opensource software licenses.

There have been some earlier attempts at software that enables RSA methods: There is an initial RSA library for MATLAB [113]. In addition, some steps of RSA analysis like pattern extraction and the calculation of simple RDMs are part of many neuroscience packages, like: pyMVPA [134], CosmoMVPA [135], MVPA-light [136]) and NeuroRA [137]. Furthermore, a Bayes approach to RSA is available in BrainIAK [138]. However, a comprehensive library that incorporates new developments was missing.

### 3.3 Other approaches to comparing representations

There are three commonly used alternative methods for comparing representations, which have some underlying relationships with RSA, but are usually viewed as distinct approaches [61]:

*Encoding models* are explicit models to predict one representation from another [139, 140]. The advantage of this approach is that it yields explicit predictions for the responses to new conditions, which can be evaluated just as any other prediction of data would be. The first drawback of this approach is the complexity of the encoding model. Even linear models become so flexible in high dimensions that even our largest datasets are insufficient to strongly constrain the parameters of the encoding model. Thus, regularization, dimensionality reduction and similar techniques are necessary to make good predictions [141, e.g.] and these seemingly minor choices can substantially change the results. Furthermore, fitting the encoding models can become computationally expensive and there is some debate about the adequate level of flexibility encoding models should have [15]. The second main drawback of encoding models is that they yield asymmetric results, i.e. prediction of a first model from a second model may work well while prediction from the first one to the second works barely above chance. This can be acceptable behavior when we predict neural data from a model, but for comparisons between models or between data sets this is a clear disadvantage [62]. Nonetheless, encoding models remain a popular and sensible option for the evaluation of models that predict brain data.

In Pattern Component Modelling [PCM, 64] the feature weights are instead treated as random variables. PCM evaluates the likelihood of the RM given the distribution of activity profiles; the response of a measurement channel to each of the conditions. In effect the RM is described by the second moment of these activity profiles. PCM excels at model selection, but requires strong assumptions on the generative model. It arguably also lacks an intuitively understandable component statistic.

In machine learning, comparisons are more usually made based on the kernel matrices of representations [63, centered kernel alignment]. Due to the strong connection between kernel and distances matrices [70], these methods can be seen as part of the same family of methods. Indeed, we implement centered kernel alignment (CKA, the main measure of this class) in our toolbox.

Beyond these main approaches there a few additional ones that do not fall into one of these categories. Two methods that differ from methods that are invariance to rotation of the neural space are the Soft Matching Distance [142] a metric sensitive to the tuning of individual neurons, and tuning reorientation [143], a measure of alignment complexity.

Which metrics are best used under what circumstances is still debated [144, 145] and researchers are actively testing and comparing methods against each other. Thus, we follow an inclusive strategy for our toolbox and will in the future keep integrating new promising metrics.

### 3.4 Conclusion

Our new *rsatoolbox* implements all state-of-the-art methods of representational similarity analysis and constitutes the open-source development project where this methodology will be further developed. It has data reading functions to import behavioral, EEG/MEG and fMRI data into structured objects. RDMs can then be computed from these and subsequently manipulated along various keys and dimensions. These RDMs can then be compared and used to evaluate models with proper statistics. Visualization functions make it easy to plot the RDMs and results. Thus, rsatoolbox gives researchers access to some of the latest methods in RSA, without a steep learning curve, while potentially making their analyses faster. By building a community project we aim for RSAtoolbox to be the standard starting point of any RSA analysis.

## Acknowledgments

Many thanks to everyone who has contributed ideas, code, documentation, or feedback to rsatoolbox. JB is supported by a UKRI BBSRC grant (BB/X008428/1) awarded to Faisal Mushtaq. We thank all the contributors to open-source software without which we could not have made rsatoolbox, in particular Python, NumPy, scipy, cython, conda-forge and many others.

## Notes

### Competing Interest Statement

The authors have declared no competing interest.

## References

[1] Allen, E. J. et al. A massive 7T fMRI dataset to bridge cognitive neuroscience and artificial intelligence. Nature neuroscience 25, 116–126 (2022).

[2] Hebart, M. N. et al. THINGS-data, a multimodal collection of large-scale datasets for investigating object representations in human brain and behavior. eLife 12 (2023).

[3] Kupers, E. R., Knapen, T., Merriam, E. P. & Kay, K. N. Principles of intensive human neuroimaging. Trends in neurosciences 47, 856–864 (2024).

[4] de Vries, S. E. J. et al. A large-scale standardized physiological survey reveals functional organization of the mouse visual cortex. Nature neuroscience 23, 138–151 (2020).

[5] International Brain Laboratory et al. Standardized and reproducible measurement of decision-making in mice. eLife 10 (2021).

[6] Papale, P., Wang, F., Self, M. W. & Roelfsema, P. R. An extensive dataset of spiking activity to reveal the syntax of the ventral stream. Neuron 113, 539–553.e5 (2025).

[7] Churchland, A. K. & Abbott, L. F. Conceptual and technical advances define a key moment for theoretical neuroscience. Nature neuroscience 19, 348–349 (2016).

[8] Kriegeskorte, N. & Douglas, P. K. Cognitive computational neuroscience. Nature neuroscience 21, 1148–1160 (2018).

[9] Naselaris, T. et al. Cognitive computational neuroscience: A new conference for an emerging discipline. Trends in cognitive sciences 22, 365–367 (2018).

[10] Krizhevsky, A., Sutskever, I. & Hinton, G. E. ImageNet classification with deep convolutional neural networks. Communications of the ACM 60, 84–90 (2017).

[11] LeCun, Y., Bengio, Y. & Hinton, G. Deep learning. Nature 521, 436–444 (2015).

[12] Hosseini, E. A. et al. Artificial neural network language models predict human brain responses to language even after a developmentally realistic amount of training. *Neurobiology of language (Cambridge*, Mass.) 5, 43–63 (2024).

[13] Dapello, J. et al. Simulating a primary visual cortex at the front of CNNs improves robustness to image perturbations. bioRxiv 2020.06.16.154542 (2020).

[14] Baker, B. et al. What makes representations “useful”? Cognitive Computational Neuroscience 2021 (2021).

[15] Kriegeskorte, N. & Diedrichsen, J. Peeling the Onion of Brain Representations. Annual review of neuroscience 42, 407–432 (2019).

[16] Thagard, P. Cognitive Science. https://plato.stanford.edu/archives/win2023/entries/cognitive-science/ (2023).

[17] Kriegeskorte, N., Mur, M. & Bandettini, P. Representational similarity analysis - connecting the branches of systems neuroscience. Frontiers in systems neuroscience 2, 4 (2008).

[18] Kriegeskorte, N. & Kievit, R. A. Representational geometry: integrating cognition, computation, and the brain. Trends in cognitive sciences 17, 401–412 (2013).

[19] Torgerson, W. S. Theory and methods of scaling (Wiley, Oxford, England, 1958).

[20] Shepard, R. N. The analysis of proximities: Multidimensional scaling with an unknown distance function. I. Psychometrika 27, 125–140 (1962).

[21] Shepard, R. N. & Chipman, S. Second-order isomorphism of internal representations: Shapes of states. Cognitive psychology 1, 1–17 (1970).

[22] Edelman, S., Grill-Spector, K., Kushnir, T. & Malach, R. Toward direct visualization of the internal shape representation space by fMRI. Psychobiology 26, 309–321 (1998).

[23] Wang, L., Uhrig, L., Jarraya, B. & Dehaene, S. Representation of numerical and sequential patterns in macaque and human brains. Current biology: CB 25, 1966–1974 (2015).

[24] Cohen, M. A. et al. Representational similarity precedes category selectivity in the developing ventral visual pathway. NeuroImage 197, 565–574 (2019).

[25] Cichy, R. M., Pantazis, D. & Oliva, A. Similarity-Based Fusion of MEG and fMRI Reveals Spatio-Temporal Dynamics in Human Cortex During Visual Object Recognition. Cerebral cortex 26, 3563–3579 (2016).

[26] Kaneshiro, B., Perreau Guimaraes, M., Kim, H.-S., Norcia, A. M. & Suppes, P. A Representational Similarity Analysis of the dynamics of object processing using single-trial EEG classification. PloS one 10, e0135697 (2015).

[27] Ejaz, N., Hamada, M. & Diedrichsen, J. Hand use predicts the structure of representations in sensorimotor cortex. Nature neuroscience 18, 1034–1040 (2015).

[28] Devereux, B. J., Clarke, A., Marouchos, A. & Tyler, L. K. Representational similarity analysis reveals commonalities and differences in the semantic processing of words and objects. The Journal of neuroscience: the official journal of the Society for Neuroscience 33, 18906–18916 (2013).

[29] Zhao, L. et al. Orthographic and phonological representations in the fusiform cortex. Cerebral cortex (New York, N.Y.: 1991) 27, 5197–5210 (2017).

[30] Giordano, B. L., McAdams, S., Zatorre, R. J., Kriegeskorte, N. & Belin, P. Abstract encoding of auditory objects in cortical activity patterns. Cerebral cortex 23, 2025–2037 (2013).

[31] Majewska, O. et al. Semantic data set construction from human clustering and spatial arrangement. Computational linguistics (Association for Computational Linguistics) 47, 69–116 (2021).

[32] Levitan, C. A. et al. Cross-cultural color-odor associations. PloS one 9, e101651 (2014).

[33] Pauli, R. & Lockwood, P. L. The computational psychiatry of antisocial behaviour and psychopathy. Neuroscience and biobehavioral reviews 145, 104995 (2023).

[34] Dobs, K., Isik, L., Pantazis, D. & Kanwisher, N. How face perception unfolds over time. Nature communications 10, 1258 (2019).

[35] Haushofer, J., Livingstone, M. S. & Kanwisher, N. Multivariate patterns in object-selective cortex dissociate perceptual and physical shape similarity. *PLoS biology* 6, e187 (2008).

[36] Schurger, A., Pereira, F., Treisman, A. & Cohen, J. D. Reproducibility distinguishes conscious from nonconscious neural representations. Science 327, 97–99 (2010).

[37] Proklova, D., Kaiser, D. & Peelen, M. V. Disentangling representations of object shape and object category in human visual cortex: The animate-inanimate distinction. Journal of cognitive neuroscience 28, 680–692 (2016).

[38] Carlson, T., Tovar, D. A., Alink, A. & Kriegeskorte, N. Representational dynamics of object vision: the first 1000 ms. Journal of vision 13 (2013).

[39] Xue, G. et al. Greater neural pattern similarity across repetitions is associated with better memory. Science 330, 97–101 (2010).

[40] Wimber, M., Alink, A., Charest, I., Kriegeskorte, N. & Anderson, M. C. Retrieval induces adaptive forgetting of competing memories via cortical pattern suppression. Nature neuroscience 18, 582–589 (2015).

[41] Kriegeskorte, N. & Diedrichsen, J. Inferring brain-computational mechanisms with models of activity measurements. Philosophical transactions of the Royal Society of London. Series B, Biological sciences 371, 20160278 (2016).

[42] Freiwald, W. A. & Tsao, D. Y. Functional compartmentalization and viewpoint generalization within the macaque face-processing system. *Science (New York*, N.Y.) 330, 845–851 (2010).

[43] Ringach, D. The geometry of masking in neural populations. Nature communications 10, 4879 (2019).

[44] Yamins, D. L. K. et al. Performance-optimized hierarchical models predict neural responses in higher visual cortex. Proceedings of the National Academy of Sciences of the United States of America 111, 8619–8624 (2014).

[45] Sadeh, S. & Clopath, C. Contribution of behavioural variability to representational drift. bioRxiv 11 (2022).

[46] Takeda, K., Abe, K., Kitazono, J. & Oizumi, M. Unsupervised alignment reveals structural commonalities and differences in neural representations of natural scenes across individuals and brain areas. *ICLR 2024 Workshop on Representational Alignment* (2024).

[47] Khaligh-Razavi, S.-M. & Kriegeskorte, N. Deep supervised, but not unsupervised, models may explain IT cortical representation. PLoS computational biology 10, e1003915 (2014).

[48] Mehrer, J., Spoerer, C. J., Kriegeskorte, T. C., Nikolaus & Kietzmann. Individual differences among deep neural network models. Nature Communications 11 (2020).

[49] Kriegeskorte, N. Deep neural networks: A new framework for modeling biological vision and brain information processing. Annual review of vision science 1, 417–446 (2015).

[50] Yamins, D. & DiCarlo, J. Using goal-driven deep learning models to understand sensory cortex. Nature neuroscience 19, 356–365 (2016).

[51] Xu, Y. & Vaziri-Pashkam, M. Limits to visual representational correspondence between convolutional neural networks and the human brain. Nature communications 12, 2065 (2021).

[52] Konkle, T. & Alvarez, G. A. A self-supervised domain-general learning framework for human ventral stream representation. Nature communications (2022).

[53] Cichy, R. M. & Kaiser, D. Deep neural networks as scientific models. Trends in cognitive sciences 23, 305–317 (2019).

[54] Storrs, K. R., Anderson, B. L. & Fleming, R. W. Unsupervised learning predicts human perception and misperception of gloss. Nature human behaviour 5, 1402– 1417 (2021).

[55] Kietzmann, T. C. et al. Recurrence is required to capture the representational dynamics of the human visual system. Proceedings of the National Academy of Sciences 116, 21854–21863 (2019).

[56] Groen, I. I., et al. Distinct contributions of functional and deep neural network features to representational similarity of scenes in human brain and behavior. eLife 7 (2018).

[57] Doerig, A. et al. The neuroconnectionist research programme. Nature reviews. Neuroscience 24, 431–450 (2023).

[58] Golan, T., et al. Deep neural networks are not a single hypothesis but a language for expressing computational hypotheses. The behavioral and brain sciences 46, e392 (2023).

[59] Kriegeskorte, N. & Wei, X.-X. Neural tuning and representational geometry. *arXiv [q-bio.NC]* (2021).

[60] Kriegeskorte, N. & Douglas, P. K. Interpreting encoding and decoding models. Current opinion in neurobiology 55, 167–179 (2019).

[61] Diedrichsen, J. & Kriegeskorte, N. Representational models: A common framework for understanding encoding, pattern-component, and representational-similarity analysis. PLoS computational biology 13, e1005508 (2017).

[62] Williams, A. H., Kunz, E., Kornblith, S. & Linderman, S. W. Generalized shape metrics on neural representations. Advances in neural information processing systems 34, 4738–4750 (2021).

[63] Kornblith, S., Norouzi, M., Lee, H. & Hinton, G. E. Similarity of neural network representations revisited. International Conference on Machine Learning abs/1905.00414, 3519–3529 (2019).

[64] Diedrichsen, J., Ridgway, G. R., Friston, K. J. & Wiestler, T. Comparing the similarity and spatial structure of neural representations: a pattern-component model. NeuroImage 55, 1665–1678 (2011).

[65] Walther, A. et al. Reliability of dissimilarity measures for multi-voxel pattern analysis. NeuroImage 137, 188–200 (2016).

[66] Bossio Botero, V. & Kriegeskorte, N. When do measured representational distances reflect the neural representational geometry (2024). In prep.

[67] Shahbazi, M., Shirali, A., Aghajan, H. & Nili, H. Using distance on the Riemannian manifold to compare representations in brain and in models. NeuroImage 239, 118271 (2021).

[68] Brown, S. & Farivar, R. The Topology of Representational Geometry. bioRxiv 2024.02.16.579506 (2024).

[69] Lin, B. & Kriegeskorte, N. The Topology and Geometry of Neural Representations. arXiv [q-bio.NC] (2023).

[70] Diedrichsen, J. et al. Comparing representational geometries using whitened unbiased-distance-matrix similarity. *Neurons, Behavior*, Data analysis, and Theory 5 (2021).

[71] Schütt, H. H., Kipnis, A. D., Diedrichsen, J. & Kriegeskorte, N. Statistical inference on representational geometries. Elife 12, e82566 (2023).

[72] Diedrichsen, J., Yokoi, A. & Arbuckle, S. A. Pattern component modeling: A flexible approach for understanding the representational structure of brain activity patterns. NeuroImage 180, 119–133 (2018).

[73] van Rossum, G. & de Boer, J. Interactively testing remote servers using the Python programming language. CWi Quarterly 4, 283–303 (1991).

[74] Brett, M. et al. nipy/nibabel: 5.2. 1. *Zenodo* (2024).

[75] Gorgolewski, K. et al. Nipype: a flexible, lightweight and extensible neuroimaging data processing framework in python. Frontiers in neuroinformatics 5, 13 (2011).

[76] Gramfort, A. et al. MEG and EEG data analysis with MNE-Python. Frontiers in neuroscience 7, 267 (2013).

[77] Esteban, O. et al. fMRIPrep: a robust preprocessing pipeline for functional MRI. Nature methods 16, 111–116 (2019).

[78] Teeters, J. L. et al. Neurodata Without Borders: Creating a common data format for neurophysiology. Neuron 88, 629–634 (2015).

[79] [79] Rübel, O., et al. NWB:N 2.0: An Accessible Data Standard for Neurophysiology. BioRxiv 523035 (2019).

[80] Rübel, O. et al. The Neurodata Without Borders ecosystem for neurophysiological data science. eLife 11, e78362 (2022).

[81] Harris, C. R. et al. Array programming with NumPy. Nature 585, 357–362 (2020).

[82] Garyfallidis, E. et al. Dipy, a library for the analysis of diffusion MRI data. Frontiers in neuroinformatics 8, 8 (2014).

[83] Abraham, A. et al. Machine learning for neuroimaging with scikit-learn. Frontiers in neuroinformatics 8, 14 (2014).

[84] Moore, G. P., Perkel, D. H. & Segundo, J. P. Statistical analysis and functional interpretation of neuronal spike data. Annual review of physiology 28, 493–522 (1966).

[85] Maimon, G. & Assad, J. A. Beyond Poisson: increased spike-time regularity across primate parietal cortex. Neuron 62, 426–440 (2009).

[86] Shinomoto, S., Shima, K. & Tanji, J. Differences in spiking patterns among cortical neurons. Neural computation 15, 2823–2842 (2003).

[87] Goris, R. L. T., Movshon, J. A. & Simoncelli, E. P. Partitioning neuronal variability. Nature neuroscience 17, 858–865 (2014).

[88] Goris, R. L. T., Coen-Cagli, R., Miller, K. D., Priebe, N. J. & Lengyel, M. Response sub-additivity and variability quenching in visual cortex. Nature reviews. Neuroscience 25, 237–252 (2024).

[89] Yu, B. M. et al. Gaussian-process factor analysis for low-dimensional single-trial analysis of neural population activity. Advances in Neural Information Processing Systems 21 (2008).

[90] Engemann, D. A. & Gramfort, A. Automated model selection in covariance estimation and spatial whitening of MEG and EEG signals. NeuroImage 108, 328–342 (2015).

[91] Pernet, C. et al. Issues and recommendations from the OHBM COBIDAS MEEG committee for reproducible EEG and MEG research. Nature neuroscience 23, 1473–1483 (2020).

[92] Alink, A., Walther, A., Krugliak, A., van den Bosch, J. J. F. & Kriegeskorte, N. Mind the drift - improving sensitivity to fMRI pattern information by accounting for temporal pattern drift. bioRxiv 032391 (2015).

[93] Worsley, K. J. & Friston, K. J. Analysis of fMRI time-series revisited–again. NeuroImage 2, 173–181 (1995).

[94] Prince, J. S. et al. Improving the accuracy of single-trial fMRI response estimates using GLMsingle. eLife 11 (2022).

[95] Mumford, J. A., Turner, B. O., Ashby, F. G. & Poldrack, R. A. NeuroImage Deconvolving BOLD activation in event-related designs for multivoxel pattern classi fi cation analyses. NeuroImage 59, 2636–2643 (2012).

[96] Friston, K. in Statistical parametric mapping: The analysis of functional brain images (eds Penny, W. D., Friston, K. J., Ashburner, J. T., Kiebel, S. J. & Nichols, T. E.) Statistical parametric mapping: The analysis of functional brain images (Academic Press, 2011).

[97] Gorgolewski, K. J. et al. The brain imaging data structure, a format for organizing and describing outputs of neuroimaging experiments. Scientific data 3, 160044 (2016).

[98] Op de Beeck, H., Wagemans, J. & Vogels, R. Inferotemporal neurons represent low-dimensional configurations of parameterized shapes. Nature neuroscience 4, 1244–1252 (2001).

[99] Kiani, R., Esteky, H., Mirpour, K. & Tanaka, K. Object category structure in response patterns of neuronal population in monkey inferior temporal cortex. Journal of neurophysiology 97, 4296–4309 (2007).

[100] Hebart, M. N., Zheng, C. Y., Pereira, F. & Baker, C. I. Revealing the multidimensional mental representations of natural objects underlying human similarity judgements. Nature human behaviour 4, 1173–1185 (2020).

[101] Zheng, C., Pereira, F., Baker, C. & Hebart, M. Revealing interpretable object representations from human behavior. International Conference on Learning Representations abs/1901.02915 (2019).

[102] Ratan Murty, N. A., Bashivan, P., Abate, A., DiCarlo, J. J. & Kanwisher, N. Computational models of category-selective brain regions enable high-throughput tests of selectivity. Nature communications 12, 5540 (2021).

[103] Golan, T., Guo, W., Schütt, H. H. & Kriegeskorte, N. Distinguishing representational geometries with controversial stimuli: Bayesian experimental design and its application to face dissimilarity judgments. arXiv [q-bio.NC] (2022).

[104] Styrnal, M., Kaniuth, P., Stoinski, L. & Hebart, M. N. What do similarity tasks actually measure? A systematic comparison of eight tasks. Workshop on concepts, actions, and objects: Functional and neural perspectives (2024).

[105] Kriegeskorte, N. & Mur, M. Inverse MDS: Inferring dissimilarity structure from multiple item arrangements. Frontiers in psychology 3, 245 (2012).

[106] Mur, M. et al. Human Object-Similarity Judgments Reflect and Transcend the Primate-IT Object Representation. Frontiers in psychology 4, 128 (2013).

[107] Dima, D. C., Tomita, T. M., Honey, C. J. & Isik, L. Social-affective features drive human representations of observed actions. eLife 11 (2022).

[108] Dima, D. C., Hebart, M. N. & Isik, L. A data-driven investigation of human action representations. Scientific reports 13, 5171 (2023).

[109] Lindh, D., Sligte, I. G., Assecondi, S., Shapiro, K. L. & Charest, I. Conscious perception of natural images is constrained by category-related visual features. Nature communications 10, 4106 (2019).

[110] Faghel-Soubeyrand, S. et al. Neural computations in prosopagnosia. Cerebral cortex (New York, N.Y.: 1991) 34, bhae211 (2024).

[111] Koenig-Robert, R., Quek, G. L., Grootswagers, T. & Varlet, M. Movement trajectories as a window into the dynamics of emerging neural representations. Scientific reports 14, 11499 (2024).

[112] Kriegeskorte, N., Formisano, E., Sorger, B. & Goebel, R. Individual faces elicit distinct response patterns in human anterior temporal cortex. Proceedings of the National Academy of Sciences of the United States of America 104, 20600–20605 (2007).

[113] Nili, H. et al. A toolbox for representational similarity analysis. PLoS computational biology 10, e1003553 (2014).

[114] Diedrichsen, J., Provost, S. & Zareamoghaddam, H. On the distribution of cross-validated Mahalanobis distances. arXiv [stat.AP] (2016).

[115] Gretton, A., Herbrich, R., Smola, A., Bousquet, O. & Scholkopf, B. Kernel methods for measuring independence. Journal of machine learning research: JMLR 6, 2075–2129 (2005).

[116] Székely, G. J., Rizzo, M. L. & Bakirov, N. K. Measuring and testing dependence by correlation of distances. Annals of statistics 35, 2769–2794 (2007).

[117] Harvey, S. E., Larsen, B. W. & Williams, A. H. Duality of Bures and shape distances with implications for comparing neural representations. arXiv [stat.ML] 11–26 (2023).

[118] Yarkoni, T. The generalizability crisis. The Behavioral and brain sciences 45 (2022).

[119] Kriegeskorte, N., Goebel, R. & Bandettini, P. Information-based functional brain mapping. Proceedings of the National Academy of Sciences of the United States of America 103, 3863–3868 (2006).

[120] Kriegeskorte, N. & Bandettini, P. Analyzing for information, not activation, to exploit high-resolution fMRI. NeuroImage 38, 649–662 (2007).

[121] Devereux, B. J., Clarke, A. & Tyler, L. K. Integrated deep visual and semantic attractor neural networks predict fMRI pattern-information along the ventral object processing pathway. Scientific reports 8, 10636 (2018).

[122] Tucciarelli, R., Wurm, M., Baccolo, E. & Lingnau, A. The representational space of observed actions. eLife 8 (2019).

[123] Oosterhof, N. N., Wiestler, T., Downing, P. E. & Diedrichsen, J. A comparison of volume-based and surface-based multi-voxel pattern analysis. NeuroImage 56, 593–600 (2011).

[124] Cichy, R. M., Pantazis, D. & Oliva, A. Resolving human object recognition in space and time. Nature neuroscience 17, 455–462 (2014).

[125] Clarke, A., Devereux, B. J., Randall, B. & Tyler, L. K. Predicting the time course of individual objects with MEG. Cerebral cortex (New York, N.Y.: 1991) 25, 3602–3612 (2015).

[126] Grootswagers, T., Wardle, S. G. & Carlson, T. A. Decoding dynamic brain patterns from evoked responses: A tutorial on multivariate pattern analysis applied to time series neuroimaging data. Journal of cognitive neuroscience 29, 677–697 (2017).

[127] Hebart, M. N., Bankson, B. B., Harel, A., Baker, C. I. & Cichy, R. M. The representational dynamics of task and object processing in humans. eLife 7 (2018).

[128] Jozwik, K. M., Kietzmann, T. C., Cichy, R. M., Kriegeskorte, N. & Mur, M. Deep neural networks and visuo-semantic models explain complementary components of human ventral-stream representational dynamics. The Journal of neuroscience: the official journal of the Society for Neuroscience 43, 1731–1741 (2023).

[129] Storrs, K. R., Khaligh-Razavi, S.-M. & Kriegeskorte, N. Noise ceiling on the crossvalidated performance of reweighted models of representational dissimilarity: Addendum to Khaligh-Razavi & Kriegeskorte (2014). bioRxiv 2020.03.23.003046 (2020).

130. Kendall, M. G. Rank correlation methods. *pp* (1948).

[131] Poldrack, R. A. & Gorgolewski, K. J. Making big data open: data sharing in neuroimaging. Nature neuroscience 17, 1510–1517 (2014).

[132] Keator, D. B. et al. Towards structured sharing of raw and derived neuroimaging data across existing resources. NeuroImage 82, 647–661 (2013).

[133] Humble, J. & Farley, D. Continuous Delivery: Reliable Software Releases through Build, Test, and Deployment Automation (Pearson Education, 2010).

[134] Hanke, M. et al. PyMVPA: A python toolbox for multivariate pattern analysis of fMRI data. Neuroinformatics 7, 37–53 (2009).

[135] Oosterhof, N. N., Connolly, A. C. & Haxby, J. V. CoSMoMVPA: Multi-Modal Multivariate Pattern Analysis of Neuroimaging Data in Matlab/GNU Octave. Frontiers in neuroinformatics 10, 27 (2016).

[136] Treder, M. S. MVPA-Light: A Classification and Regression Toolbox for Multi-Dimensional Data. Frontiers in neuroscience 14, 289 (2020).

[137] Lu, Z. & Ku, Y. NeuroRA: A python toolbox of representational analysis from multi-modal neural data. Frontiers in neuroinformatics 14, 563669 (2020).

[138] Cai, M. B., Schuck, N. W., Pillow, J. W. & Niv, Y. Representational structure or task structure? Bias in neural representational similarity analysis and a Bayesian method for reducing bias. PLoS computational biology 15, e1006299 (2019).

[139] van Gerven, M. A. J. A primer on encoding models in sensory neuroscience. Journal of mathematical psychology 76, 172–183 (2017).

[140] Naselaris, T., Kay, K. N., Nishimoto, S. & Gallant, J. L. Encoding and decoding in fMRI. NeuroImage 56, 400–410 (2011).

[141] Conwell, C., Prince, J. S., Kay, K. N., Alvarez, G. A. & Konkle, T. A large-scale examination of inductive biases shaping high-level visual representation in brains and machines. Nature communications 15, 9383 (2024).

[142] Khosla, M. & Williams, A. H. Soft Matching Distance: A metric on neural representations that captures single-neuron tuning. Proceedings of UniReps: the First Workshop on Unifying Representations in Neural Models 326–341 (2024).

[143] Prince, J. S., Conwell, C., Alvarez, G. A. & Konkle, T. A case for sparse positive alignment of neural systems. *ICLR 2024 Workshop on Representational Alignment* (2024).

[144] Conwell, C. et al. Battle of the Brain-Model-Mapping Metrics. Cognitive Computational Neuroscience 2024 (2024).

[145] Feather, J., Leclerc, G., Mądry, A. & McDermott, J. H. Model metamers reveal divergent invariances between biological and artificial neural networks. Nature neuroscience 26, 2017–2034 (2023).

